# Creation of de novo cryptic splicing for ALS/FTD precision medicine

**DOI:** 10.1101/2023.11.15.565967

**Authors:** Oscar G. Wilkins, Max Z.Y.J. Chien, Josette J. Wlaschin, Maria Pisliakova, David Thompson, Holly Digby, Rebecca L. Simkin, Juan Antinao Diaz, Puja R. Mehta, Matthew J. Keuss, Matteo Zanovello, Anna-Leigh Brown, Peter Harley, Annalucia Darbey, Rajvinder Karda, Elizabeth M.C. Fisher, Tom J. Cunningham, Claire E. Le Pichon, Jernej Ule, Pietro Fratta

**Affiliations:** UCL Queen Square Motor Neuron Disease Centre, Department of Neuromuscular Diseases, UCL Queen Square Institute of Neurology, UCL; London, WC1N 3BG, UK; The Francis Crick Institute; London, NW1 1AT, UK; Eunice Kennedy Shriver National Institute of Child Health and Human Development, National Institutes of Health; Bethesda, MD 20892, USA; Mammalian Genetics Unit, MRC Harwell Institute; Oxfordshire, OX11 0RD, UK; UK Dementia Research Institute at King’s College London, Maurice Wohl Clinical Neuroscience Institute; London, SE5 9RX, UK; EGA-Institute for Women’s Health, University College London; London, WC1E 6HX, UK

## Abstract

A system enabling the expression of therapeutic proteins specifically in diseased cells would be transformative, providing greatly increased safety and the possibility of pre-emptive treatment. Here we describe “TDP-REG”, a precision medicine approach primarily for amyotrophic lateral sclerosis (ALS) and frontotemporal dementia (FTD), which exploits the cryptic splicing events that occur in cells with TDP-43 loss-of-function (TDP-LOF) in order to drive expression specifically in diseased cells. In addition to modifying existing cryptic exons for this purpose, we develop a deep-learning-powered algorithm for generating customisable cryptic splicing events, which can be embedded within virtually any coding sequence. By placing part of a coding sequence within a novel cryptic exon, we tightly couple protein expression to TDP-LOF. Protein expression is activated by TDP-LOF *in vitro* and *in vivo*, including TDP-LOF induced by cytoplasmic TDP-43 aggregation. In addition to generating a variety of fluorescent and luminescent reporters, we use this system to perform TDP-LOF-dependent genomic prime editing to ablate the *UNC13A* cryptic donor splice site. Furthermore, we design a panel of tightly gated, autoregulating vectors encoding a TDP-43/Raver1 fusion protein, which rescue key pathological cryptic splicing events. In summary, we combine deep-learning and rational design to create sophisticated splicing sensors, resulting in a platform that provides far safer therapeutics for neurodegeneration, potentially even enabling preemptive treatment of at-risk individuals.

**One-Sentence Summary:** We engineer TDP-43-regulated cryptic exons, enabling exceptionally precise activation of gene therapies in diseased neurons.

## Main Text

Amyotrophic lateral sclerosis (ALS) is a devastating and incurable neurodegenerative disease. In 97% of cases, there is pronounced formation of neuronal cytoplasmic aggregates of the RNA-binding protein TDP-43 (*1*). Further, such pathology is found in ∼45% of frontotemporal dementia (FTD) cases, and is also observed in limbic-predominant age-related TDP-43 encephalopathy (LATE) and Alzheimer’s disease, suggesting TDP-43 mediated RNA dysregulation is a critical element in many neurodegenerative diseases and therefore a promising clinical target (*2*, *3*).

TDP-43 plays a crucial role in the regulation of splicing, protecting the transcriptome from the inclusion of toxic “cryptic exons” (CEs), which are rarely detected in healthy cells, but become prominently expressed upon TDP-LOF; CEs typically introduce premature termination codons (PTCs) into transcripts, preventing the expression of crucial proteins including STMN2 and UNC13A (*4–8*). We and others previously showed that genetic modulation of even a single specific cryptic exon can influence disease progression, meaning the stage in which cryptic exons are expressed is within the therapeutic window (*7*, *8*). Numerous preclinical studies aim to reduce TDP-43-associated toxicity, for example via blocking cryptic exon inclusion using antisense oligonucleotides (ASOs) or transgenes, or by targeting the aggregation process itself (*4*, *9–11*).

A major barrier to bringing gene therapy approaches to the clinic is the lack of methods to tightly regulate transgene expression. Only a small fraction of motor neurons, let alone all neurons, display clear pathology at any given time in patients (*12*). Therefore, a conventional targeting approach (for example, combining an AAV serotype with CNS-tropism with a neuronal promoter) would result in expression of the therapeutic transgene in a vast number of non-degenerating cells. Such expression is not merely unnecessary, but could actually worsen prognosis by interfering with the homeostasis of otherwise healthy cells, especially given the permanent nature of these approaches. Such concerns recently led to the development of therapeutics with activity-dependent transcriptional promoters for epilepsy; however, it is unclear how a similar promoter-based approach could be applied to neurodegeneration (*13*).

Here, we leverage our understanding of TDP-43’s regulation of splicing together with deep-learning-based splicing prediction tools to generate gene therapy vectors featuring novel cryptic splice sites. Crucially, whereas pathological cryptic splicing blocks protein expression, our vectors explicitly require the use of the cryptic splice site(s) to express the encoded transgene. This approach limits expression of transgenes to cells with TDP-LOF, which co-occurs with TDP-43 aggregation in patients, thus greatly reducing the risk of off-target side-effects. In addition to demonstrating highly effective TDP-LOF-dependent expression *in vitro* and *in vivo*, we apply this approach to two candidate ALS/FTD gene therapies, paving the way towards safer and more efficacious treatments for these devastating diseases.

### TDP-REGv1: Modification of ALS/FTD cryptic exons for TDP-43-regulated expression vectors

Numerous TDP-43-regulated CEs have been reproducibly detected in cellular and animal models, showing high specificity for TDP-LOF; furthermore, a number of these, including CEs in *STMN2*, *UNC13A*, *HDGFL2* and *AARS1*, have been found specifically within neurons with TDP-43 pathology in post-mortem ALS and FTD samples (*4–8*, *14*). We reasoned that if configured in a suitable manner, a cryptic exon with these features could be used to regulate expression of a therapeutic or reporter transgene (**Fig. 1A**). We identified the cryptic exon in the *AARS1* gene (hg38: chr16:70272796-70272882) as the most suitable candidate owing to its reproducible detection, its lack of stop codons in at least one frame, and its short, defined TDP-43 binding motif (**Fig. 1B; fig. S1)**. To make it suitable for use within a frame-shifting “minigene”, we added an extra residue to the cryptic exon to cause frame-shifting, and removed sections of the flanking introns that were likely unnecessary for regulation by TDP-43 (**Fig. 1B**). We then placed this sequence between an upstream start codon and downstream transgene such that only when the cryptic exon was included would the start codon be in-frame with the transgene (**Fig. 1B**). We included a P2A “self-cleavage” site so that the upstream peptide encoded by the CE is released separately from the main protein (**Fig. 1B**). Furthermore, we added a constitutively-spliced intron derived from *RPS24* to promote nonsense-mediated decay (NMD) in the event that a PTC is encountered.

**Fig. 1.**
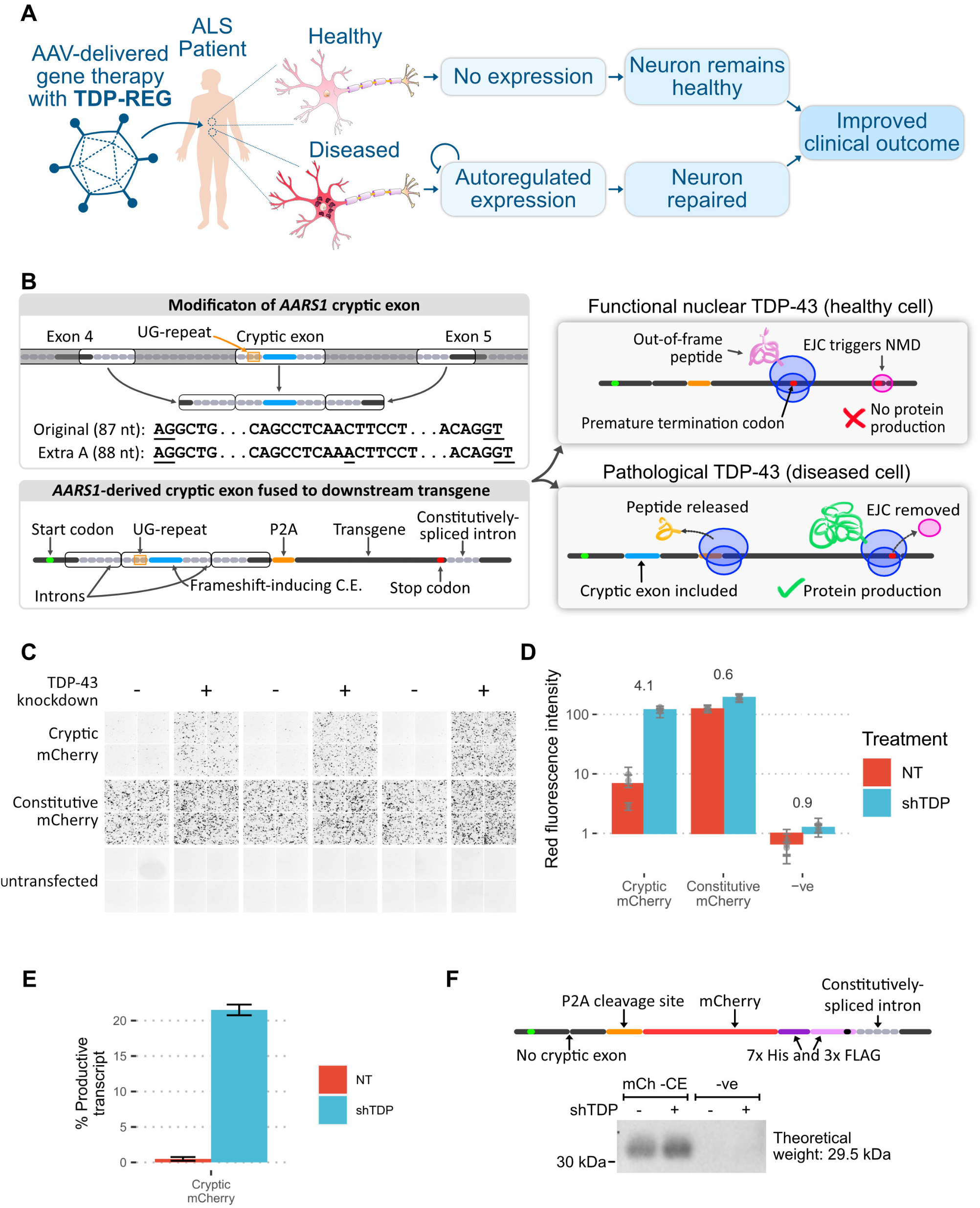
Regulatory upstream CE controls downstream transgene expression (TDP-REGv1). **A:** Schematic showing the intended function and purpose of this work. **B:** Schematic of TDP-REGv1. *Top left:* the genomic locus surrounding the human *AARS1* CE region was modified, with the middle parts of the introns removed to reduce size, and an extra adenosine added to the CE sequence to enable frame-shifting. The AG and GT splice sites of the CE are underlined. *Bottom left:* the modified sequence from part A was incorporated into a “minigene”. *Top right*: when the CE is excluded, the start codon is out-of-frame with the transgene, triggering nonsense-mediated decay (NMD) due to the downstream exon junction complex (EJC). *Bottom right:* when the CE is included, the transgene is in-frame with the start codon, resulting in protein expression. **C**: Fluorescence microscopy images (red channel) showing SK-N-BE(2) cells with (shTDP) or without (NT = “not treated”) TDP-43 knockdown, transfected with a vector fusing the upstream *AARS1-*derived sequence to mCherry (“cryptic mCherry”) or a constitutive vector. **D**: Quantification of the images in Part C; numbers show log2-fold-change in TDP-43 knockdown cells; each dot shows the average of one well (three wells per condition), with error bars showing the standard deviation within each well (four per well). **E**: Summary of Nanopore sequencing results for the cells in Part C; error bars show standard deviation across three replicates. **F**: Top: schematic of mCh -CE vector, without the CE but with a downstream His/Tri-FLAG dual tag to enable sensitive detection. Bottom: Western blot of cells transduced with the above vector (“mCh-CE”) or a completely different vector (“-ve” - a Prime Editing vector). Samples were enriched with a His-tag pulldown, then blotted with an anti-FLAG antibody.

To test if this system restricted expression to cells with TDP-LOF, we used mCherry as the transgene. This reporter construct, along with a positive control in which the cryptic exon sequence is constitutively expressed, was transfected into SK-N-BE(2) cells (a human neuroblastoma cell line) with doxycycline-inducible TDP-43 knockdown. A >16-fold increase in mCherry expression was detected in cells with TDP-43 knockdown when using the cryptic construct (**Fig. 1C-D**). In agreement with this, Nanopore sequencing revealed a >43-fold increase in cryptic exon inclusion upon TDP-43 knockdown (**Fig. 1E**). We noted that in cells with normal TDP-43 levels, mild leaky expression of mCherry protein was detectable, even though cryptic exon inclusion was less than 0.5%, suggesting a mechanism of protein translation that circumvents the presence of the upstream frameshift. Using western blotting, we observed low-level leaky expression even in a vector in which the cryptic exon is deleted, potentially due to use of alternative transcriptional start sites or leaky ribosomal scanning (**Fig. 1F**).

### TDP-REGv2: Creation of de novo cryptic splicing sequences

We reasoned that if new CEs could be created within the transgene-encoding region of the vector, this would remove any risk of leaky expression when the CE is spliced out because the full, uninterrupted transgene coding sequence would not be present in the mature mRNA. Further, this approach would avoid expression of unwanted upstream peptides and reduce the vector size. In pilot tests, we found that the *AARS1-*derived cryptic exon could be replaced with other sequences, and that the cryptic exon strength of these sequences correlated with SpliceAI splicing predictions (**fig. S2A-D**) (*15*). Encouraged by this, we built a deep-learning-driven *in silico* directed evolution algorithm, *SpliceNouveau*, to design *de novo* TDP-43-regulated cryptic cassette exons or single cryptic splice sites (**Fig. 2A-C)**. TDP-43 preferentially binds to stretches of UG-repeats (*16*, *17*). We therefore programmed *SpliceNouveau* to incorporate different types of UG-rich regions into our designs, including upstream UG-repeats (as in *AARS1*), downstream UG-repeats (as in *Sars*), upstream and downstream repeats (as in *Smg5*) or UG-rich regions without extended UG-repeats (as in *UNC13A*) (**fig. S3A**).

**Fig. 2.**
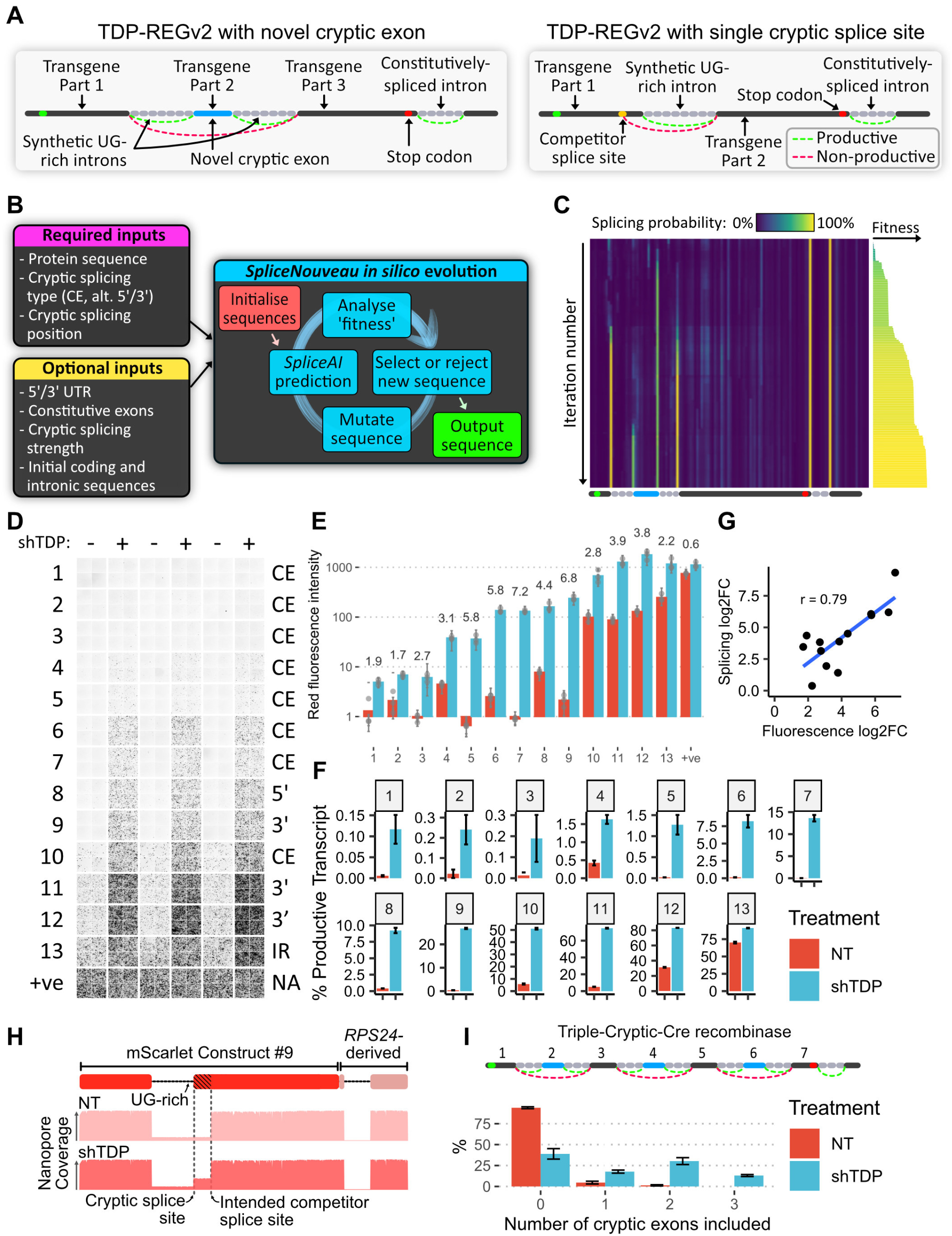
AI-guided design of novel cryptic splicing events (TDP-REGv2): **A:** Diagrams of internal cryptic exon (left) and single intron (right) designs; for the single intron design, an alternative 5’ splicing design is shown. **B:** Schematic of the *SpliceNouveau* algorithm for designing new cryptic splicing-dependent expression vectors. **C:** Heatmap showing the *in silico* evolution trajectory for an example internal cryptic exon design (below), with associated “fitness” of each (right), which was initialised with a constitutively spliced intron from *RPS24* at the 3’ end. As the iteration number increases, splice sites are “evolved” at the desired positions, and off-target splice sites are depleted. The splice sites flanking the cryptic exon are weaker than the constitutive splice sites, as specified by the user. **D:** Fluorescence microscopy images showing mScarlet expression for 13 constructs generated by *SpliceNouveau* and a positive control. Each replicate consists of four images taken from different parts of the well. **E:** Quantification of D; each dot represents the average of the four images for each replicate; error bars show the standard deviation of images in each well. **F:** Percentage of productive transcripts (i.e. transcripts which are predicted to produce full-length mScarlet), as determined by Nanopore sequencing; error bars show standard deviation across three replicates. **G:** Correlation of fold changes in fluorescence and productive isoform fraction; Pearson correlation shown. **H**: Diagram of mScarlet construct 9 (top); representative Nanopore pileup with and without TDP-43 knockdown. **I:** Schematic of construct encoding Cre recombinase, split across seven exons; exons 2, 4 and 6 are flanked by UG-rich regions (top); number of cryptic exons included in each transcript without and with shTDP, assessed by Nanopore sequencing (bottom); error bars show standard deviation across three replicates.

To create single-intron vectors, which can be smaller than those based on CEs flanked by two introns, we modified our algorithm to design candidate alternative 5’ or alternative 3’ splice sites, which compete with the designed cryptic splice site to aid TDP-43-mediated repression (**Fig. 2A**). In each case, the single intron was heavily enriched for TDP-43 binding sites; for alternative 3’ splice sites, the algorithm optimised part of the coding sequence to produce a polypyrimidine rich region. Using *SpliceNouveau*, we also attempted to design variants exhibiting TDP-43-regulated intron retention (IR) by specifying lower SpliceAI scores (i.e. weaker splice sites) combined with a single UG-rich intron.

Of the 27 designs that were tested in HEK293 cells (immortalised human embryonic kidney cells) which explored the full range of possible design parameters, 13 gave a significant increase in expression upon TDP-43 knockdown (**Fig. 2D-H).** Many constructs achieved greater dynamic range than our initial *AARS1*-based design, with two constructs achieving >100-fold increase in expression upon TDP-43 knockdown. The maximal expression level varied greatly across constructs, from >100-fold less to greater than the positive control, thus demonstrating that expression can be fine-tuned (**Fig. 2E**).

Seven of the successful designs were further tested in an SH-SY5Y cell line (human neuroblastoma cell line) with inducible TDP-43 knockdown (*7*), which yielded very similar results, demonstrating that the splicing regulation of these vectors is robust across cellular contexts (**fig. S3B-C**). In order to demonstrate that the splicing regulation was due specifically to TDP-43 depletion, we co-transfected a selection of the above constructs into SK-N-BE(2) cells with TDP-43 knockdown, together with either functional TDP-43/Raver1 fusion protein, which has been shown to rescue TDP-LOF, or an inactive TDP-43/Raver1 fusion protein that lacks ability to bind RNA (“2FL”) (*4*, *16*). Cells co-transfected with the 2FL mutant exhibited greatly increased fluorescence compared to cells co-transfected with functional TDP-43/Raver1, confirming the key role of TDP-43 in regulating the splicing of our TDP-REG constructs (**fig. S3D).**

### Multiple cryptic exons further increase specificity

Several TDP-REGv2 constructs showed remarkable specificity, but there could be therapeutic approaches, e.g. involving genome editing, where leaky expression is extremely undesirable. We therefore examined whether multiple cryptic exons could be included within the same construct, further reducing the risk of leaky expression. We used *SpliceNouveau* to design a construct encoding Cre recombinase enzyme split across seven exons, three of which were cryptic (**Fig. 2I)**. Nanopore sequencing revealed that in untreated cells, a small proportion of transcripts featured one or two cryptic exons, but <0.05% of transcripts featured all three cryptic exons, whereas shTDP cells expressed all three cryptic exons >10% of the time (**Fig. 2I)**; using multiple cryptic exons thus further decreased the risk of leaky expression.

### TDP-REG is activated by TDP-LOF *in vivo*

Having demonstrated our constructs are functional in multiple cell lines, we examined whether they are also functional *in vivo* in mammalian spinal cord, the main target of ALS therapeutic approaches. We injected AAVs (PHP.eB serotype with hSynapsin promoter) containing TDP-REGv1 mCherry or TDP-REGv2 mScarlet (construct #7) into TDP-43 conditional knockout (cKO) mice (TDP-43^Fl/Fl^;Chat-Cre^+/wt^), where TDP-43 loss was driven by a ChAT Cre driver line and therefore directed, within the spinal cord, to motor neurons and other spinal cholinergic cells.

We observed striking expression of mCherry or mScarlet in the motor neurons of these mice, with at least 50% of identified motor neurons having clear expression in all but one case for each construct (**Fig. 3A, 3C, 3E; fig. S4A)**. By contrast, control mice (TDP-43^Fl/wt^;Chat-Cre^+/wt^) displayed very little expression of mCherry, with between 0% and 2% of identified motor neurons having detectable mCherry or mScarlet expression (**Fig. 3B, 3D, 3E; fig. S4A)**. A positive control mScarlet AAV (i.e. without TDP-REG) showed no such specificity when injected into control mice, instead strongly expressing mScarlet in the majority of TDP-43-positive motor neurons (**Fig. 3E, fig. S4B)**.

**Fig. 3.**
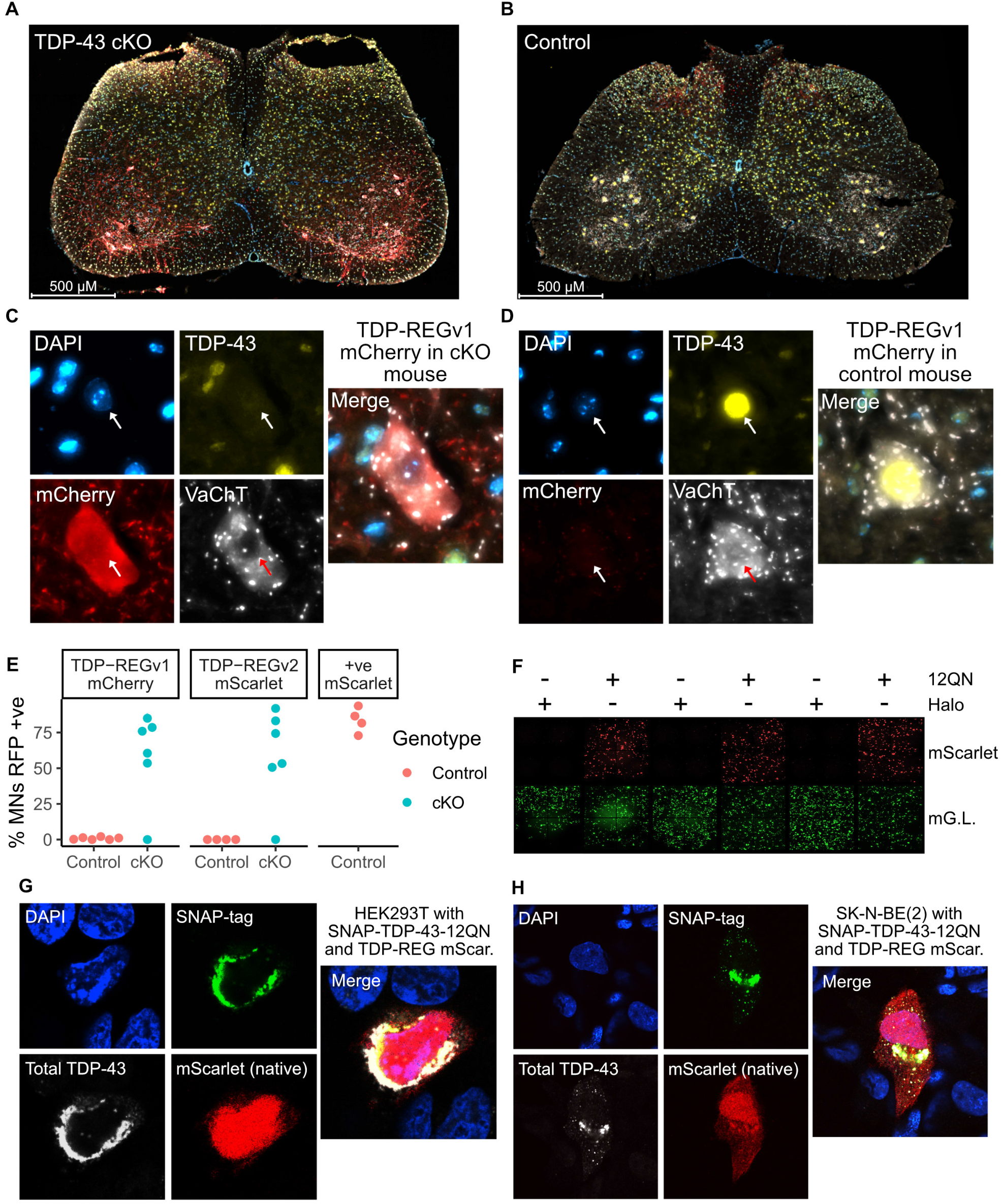
Functionality of TDP-REG in different biological contexts: **A:** Spinal cord of a TDP-43 cKO mouse injected with TDP-REGv1 mCherry AAV (blue = DAPI, yellow = TDP-43, red = mCherry, white = VAChT). **B:** Equivalent to part A, but with control mouse. **C** and **D:** Magnified representative motor neurons from figures in parts **A** and **B** respectively. **E:** Manual quantification of the percentage of motor neurons (MNs) with clear mCherry (TDP-REGv1 mCherry), mScarlet (TDP-REGv2 mScarlet #7 - see fig. S4A) or positive control mScarlet for cKO and control mice (only control mice for positive control AAV); N = 4-6 per condition. **F:** Fluorescence microscopy imaging of HEK293T cells transduced with an mScarlet TDP-REG reporter, a constitutive mGreenLantern vector, and either SNAP-TDP-43-12QN or Halo-tag expression vector. **G** and **H**: Confocal microscopy of HEK293T and SK-N-BE(2) cells, respectively, co-transfected with a TDP-REG mScarlet reporter (#7) and SNAP-TDP-43-12QN.

### TDP-43 aggregation activates TDP-REG expression

In the above experiments, TDP-43 pathology was simulated by silencing TDP-43 expression. Next, we tested whether our vectors could be activated by TDP-43 cytoplasmic aggregation, which would more accurately simulate the disease process. We co-transfected our reporters (and an mGreenLantern transfection control) into HEK293T cells with an aggregation-prone version of TDP-43 in which the Q/N-rich domain is repeated twelve times (SNAP-TDP-43-12QN), or with a vector expressing Halo-tag as a negative control (*18*). We observed strong expression of mScarlet induced by SNAP-TDP-43-12QN but not by Halo-tag (**Fig. 3F**). Using confocal microscopy, we confirmed that, in a subset of cells, SNAP-TDP-43-12QN was highly expressed and heavily enriched in the cytoplasm, activating TDP-REG mScarlet (construct #7) expression (**Fig. 3G-H).** TDP-REG is functional *in vivo* and is activated by TDP-43 aggregation, strongly suggesting this approach will function within ALS/FTD patients.

### Application of TDP-REG to biomarker expression

Next, we sought to apply TDP-REG to translationally-relevant systems. Although TDP-LOF is an established key feature of ALS progression, there is a lack of tools to detect it in cell and animal models of disease, thus limiting preclinical studies. Our fluorescent reporters described above offer a solution to detect TDP-LOF in cell studies though live imaging or FACS. We also decided to generate a sensitive luminescent biomarker that could potentially be adapted to monitor TDP-LOF in organoid and animal models. We fused the Gaussia Princeps luciferase (Gluc) sequence downstream of the *AARS1-*based minigene described above (TDP-REGv1), and in parallel also used *SpliceNouveau* to create five vectors encoding Gluc with an internal, synthetic cryptic exon (TDP-REGv2). Nanopore sequencing revealed that all of these vectors featured increased CE PSI (percentage spliced in) upon TDP-43 knockdown (**fig. S5A).** Both the TDP-REGv1 vector and the best TDP-REGv2 vectors featured high CE inclusion upon TDP-43 knockdown with minimal leakiness, but the best-performing TDP-REGv2 vector had better dynamic range (**Fig. 4A-B**): it led to >200-fold increase in Gluc expression upon TDP-43 knockdown in SK-N-BE(2) cells, and had negligible expression in cells without TDP-43 knockdown (**Fig. 4C**).

**Fig. 4.**
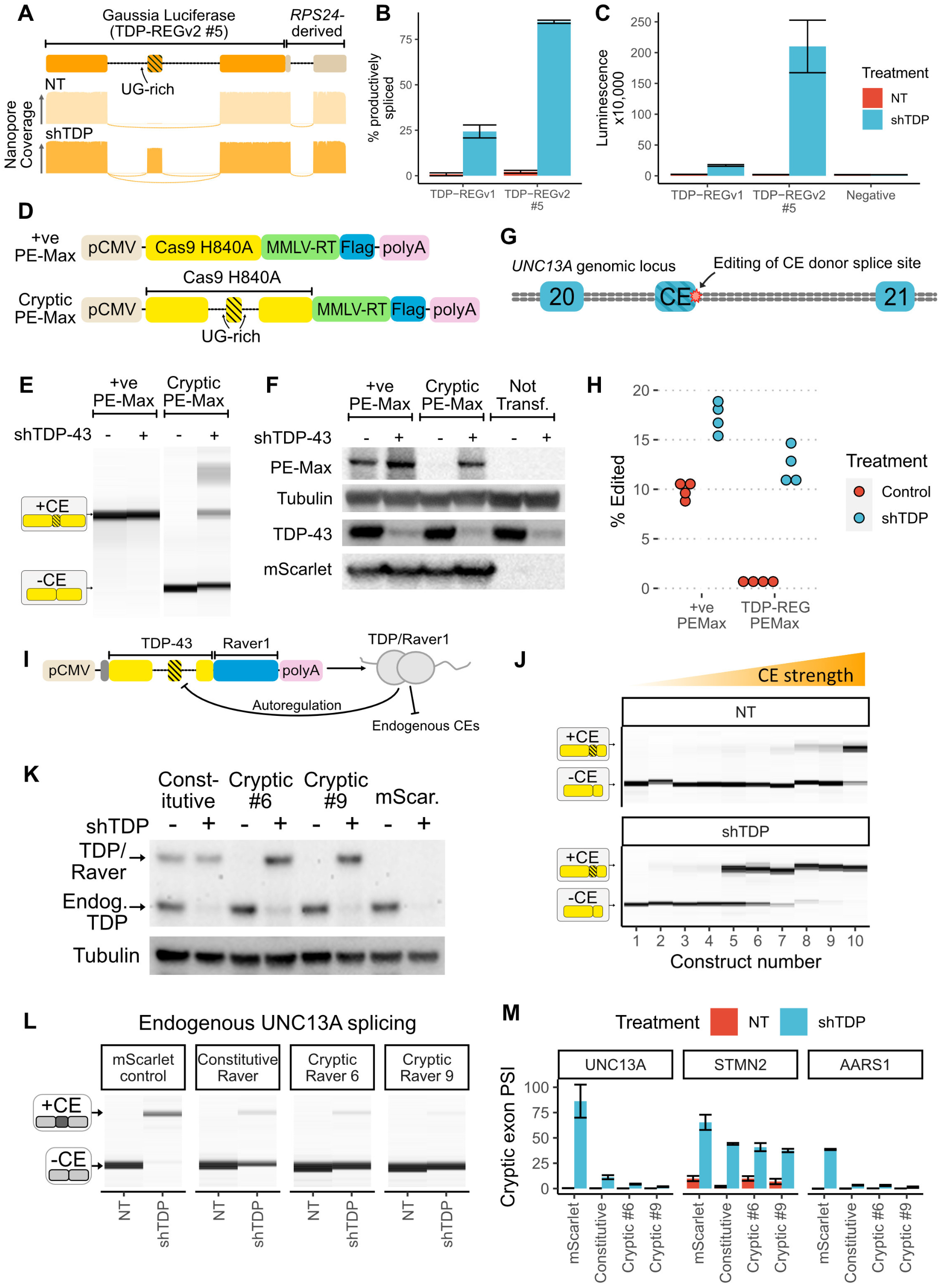
Biosensors, genome editing and splicing rescue: **A:** Schematic of TDP-REGv2 Gluc vector #5 (top); representative Nanopore pileups with (shTDP) and without (NT=”not treated” with doxycycline) TDP-43 knockdown. **B**: Quantification of Nanopore data for Gluc TDP-REGv1 and TDP-REGv2 #5 for SK-N-BE(2) cells with and without TDP-43 knockdown; error bars show standard deviation across four replicates. **C:** Quantification of luminescence from supernatant of cells used for Part B; error bars show standard deviation across four replicates. **D**: Schematic of constitutive and cryptic (TDP-REGv2) Prime Editing vector (based on Addgene PE-Max vector). **E**: Capillary electrophoresis results from RT-PCR of cryptic or constitutive PE-Max vector transfected into SK-N-BE(2) cells with or without TDP-43 knockdown. **F:** Western blot of FLAG-tagged PE-Max vectors (co-transfected with FLAG-tagged mScarlet) in SK-N-BE(2) cells with or without TDP-43 knockdown. **G:** Diagram of *UNC13A* genomic locus surrounding the *UNC13A* CE between exons 20 and 21; the position of genome editing is shown. **H**: Quantification of intended editing of *UNC13A* in SK-N-BE(2) cells with either vector, +/- TDP-43 knockdown, assessed by targeted Nanopore sequencing. **I:** Schematic of TDP-43/Raver1 with an internal cryptic exon. The design features an additional dual N-terminal NLS; the constitutive vector encodes the same amino acid sequence, but with the introns removed. **J:** RT-PCR analysis of ten constructs designed (as in part A), using the 2FL mutation to block auto-repression of the cryptic exon; constructs were transfected into SK-N-BE(2) cells without (NT) or with (shTDP) doxycycline-inducible TDP-43 knockdown. **K:** Western blot of SK-N-BE(2) cells stably expressing constitutive or cryptic (vectors 6 and 9) TDP-43/Raver1, or mScarlet, with or without endogenous TDP-43 knockdown. Endogenous TDP-43 and TDP-43/Raver1 are labelled in the anti-TDP-43 blot; tubulin loading control is shown below. All lines are polyclonal piggyBac lines; three polyclonal lines were made per construct with consistent results. **L:** RT-PCR analysis of endogenous UNC13A cryptic splicing for the same lines as Part K, showing results from a single replicate. **M:** Quantification of UNC13A, STMN2 and AARS1 cryptic splicing, using all three polyclonal replicates; error bars show standard deviation across three replicates (one replicate for mScarlet control “NT” was excluded because the RNA failed QC).

### TDP-REG tightly regulates genome editing

We explored the possibility of minimising the toxicity associated with TDP-LOF by performing genome editing to remove cryptic splice sites. The Prime Editing (PE) approach, which avoids the need for highly mutagenic double-stranded breaks, can be adapted for high-efficiency *in vivo* editing of the central nervous system (CNS) (*19*, *20*). However, even PE can lead to unwanted editing events that could potentially be toxic (*21*). Using *SpliceNouveau*, we therefore designed a TDP-REGv2 vector encoding an optimised PE construct (“PE-Max”) where part of the Cas9 sequence was encoded by a novel CE (**Fig. 4D**) (*22*). Via RT-PCR and western blotting, we confirmed that the CE had near-undetectable leaky expression (**Fig. 4E-F**).

We then designed a pegRNA to edit the donor splice site of the *UNC13A* cryptic exon (**Fig. 4G**) (*7*, *8*), combined with a nicking sgRNA to improve editing efficiency (*19*). We tested editing efficiency with or without TDP-43 knockdown in SK-N-BE(2) cells. The constitutive PE-Max vector led to editing regardless of TDP-43 knockdown, whereas the CE-containing vector tightly restricted editing to cells with TDP-43 knockdown (**Fig. 4H**). Thus, TDP-REG can ensure that genome editing occurs only in cells where it is beneficial, minimising the potential for off-target effects when delivered to all cells.

### TDP-REG can enable autoregulated splicing rescue

Finally, we used our system to directly rescue TDP-LOF. Fusions of TDP-43 to a less aggregation-prone splicing repressor domain (e.g. from *RAVER1*) can partially rescue TDP-43 splicing function (*4*). However, even small imbalances in TDP-43 expression can cause significant toxicity, and therefore in spite of its lesser aggregation, generalised expression of TDP-43/Raver1 fusions is undesirable (*23*). We therefore used *SpliceNouveau* to design TDP-REGv2 constructs encoding TDP-43 fused to the *Raver1* splicing repressor domain (TDP-REGv2:Raver1) featuring an internal CE (**Fig. 4I**). To ensure that expression of the synthetic CE occurred upon even a mild decrease in TDP-43 function, thus enabling rescue at the first signs of TDP-43 pathology, we specified high SpliceAI scores (60-100%) for the CE splice sites and used short UG-repeats to limit the binding affinity of the introns to TDP-43.

Using RT-PCR, we analysed the splicing of our constructs, with or without endogenous TDP-43 knockdown. TDP-43/Raver1 fusion protein mimics TDP-43’s splicing repressor function and would therefore be expected to autoregulate by inhibiting expression of the synthetic CEs in these constructs. Whilst this is a great advantage therapeutically, as it would prevent overexpression of TDP-43/Raver1, it could also mask the responsiveness of our synthetic CEs to TDP-LOF during these initial tests. We therefore used a 2FL mutant during these initial screens (*16*). All but one construct displayed detectable CE expression upon TDP-43 knockdown, with six showing high expression; presumably due to the use of high SpliceAI scores and relatively short UG-rich regions, some vectors had detectable leaky CE expression (**Fig. 4J**).

Using the piggyBac system, we generated polyclonal SK-N-BE(2) lines expressing TDP-43/Raver1 fusion in triplicate, either as a constitutive version or with an internal CE from constructs #6 or #9, or mScarlet. In this case, the wild-type TDP-43 sequence was used to enable both repression of cryptic splicing and autoregulation. Whereas the constitutive vector expressed TDP-43/Raver1 regardless of endogenous TDP-43 levels, vectors featuring an internal CE only expressed detectable TDP-43/Raver1 upon knockdown of endogenous TDP-43 (**Fig. 4K; fig. S5B**).

We then tested whether *TDP-REGv2:Raver1* vectors were able to repress the splicing of endogenous CEs upon TDP-43 knockdown by analysing the levels of several well-characterised CEs (**Fig. 4L-M**). Indeed, expression of TDP-43/Raver1 constructs, either constitutively spliced or with an internal CE, reduced *UNC13A* and *AARS1* CEs by over 85%, and the *STMN2* CE by approximately 40%.

Interestingly, higher levels of TDP-43/Raver1 expression, and more efficient repression of endogenous CEs, were observed upon TDP-43 knockdown for cryptic vectors (especially #9) than for the constitutive vector (**Fig. 4K-M**). We propose this is due to toxicity of TDP-43/Raver1 when expressed in cells with functional endogenous TDP-43, leading to a selection pressure against cells constitutively expressing high levels of TDP-43/Raver1. Indeed, a strong growth advantage was observed for cells expressing TDP-43/Raver1 regulated by TDP-REG compared with those expressing constitutive TDP-43/Raver1, which severely impacted growth in a dose-dependent manner (**fig. S6)**. Thus, *TDP-REGv2:Raver1* vectors can rescue endogenous CEs whilst avoiding toxicity due to constitutive TDP-43/Raver1 protein expression.

## Discussion

There is no shortage of promising new ideas for gene therapies to treat neurodegenerative disorders. However, there are major hurdles that must be overcome to translate these to the clinic. Gene therapy is typically based on a single delivery approach, which results in permanent expression throughout the patient’s lifetime, thus avoiding the need for repeated invasive delivery. However, such long-term, widespread expression can lead to significant toxicity (*24*). To mitigate this, in neurodegenerative diseases, one would ideally only express gene therapies in cells showing signs of disease, such as TDP-LOF in ALS/FTD. Crucially, this occurs only in a small proportion of cells: specific brain regions are affected in ALS and FTD, and even within the strongly affected areas, <10% of neurons show signs of TDP-43 pathology, even at the end stage of the disease (*12*). Although one can employ AAV capsids that show preference for specific tissues, and promoters can be used to limit expression to specific cell types, additional systems are needed to target expression to the small proportion of cells affected by disease pathogenesis.

Here, we describe TDP-REG, which enables disease-specific spatial and temporal regulation of gene therapy expression by exploiting the predictable splicing changes induced by TDP-LOF. Given that TDP-LOF is a hallmark of ALS, FTD, and other common neurodegenerative diseases including Alzheimer’s and LATE, TDP-REG could have broad utility in the treatment of these disorders. In at-risk individuals carrying causal genetic variants of ALS, the spatial and temporal specificity of TDP-REG could allow the therapeutics to be delivered at the pre-symptomatic stage, lying dormant until the very first stages of TDP-43 pathology are detected. Furthermore, TDP-REG can be used during the preclinical phase of drug development as a real-time readout for TDP-43 pathology in cells or even live animals.

We designed two “flavours” of TDP-REG. TDP-REGv1 involves the fusion of a transgene to an upstream regulatory module, featuring a modified CE sequence derived from an existing human gene. This approach has several benefits: as a modular design, different transgenes can be controlled by the same upstream regulatory module. Furthermore, TDP-REGv1 is based on a well-validated endogenous CE which is detected in patients in a disease-specific manner.

In contrast, TDP-REGv2 uses synthetic splicing sensors that are embedded within the transgene sequence itself. This addresses several limitations with TDP-REGv1 and previous approaches (*25*): it removes the need for a lengthy upstream regulatory region, thus aiding packaging of large transgenes into AAV vectors; it prevents an upstream, unwanted peptide being expressed and released into the cell; and it is highly tunable, meaning that splicing characteristics can be optimised for the specific transgene being delivered. For example, with a gene editing enzyme one may opt for extremely tight gating to prevent the risk of DNA damage in non-target cells, whereas for a chaperone one may prefer less tight gating if it enables higher maximal expression. Strikingly, we found that our most potent vectors had higher maximal expression levels than matched positive control vectors. This may be due to splicing promoting higher expression levels by avoiding silencing from the recently-described HUSH complex (*26*).

Splicing regulation is highly complex and difficult to predict with conventional algorithms. During the development of TDP-REGv2, we created an algorithm, *SpliceNouveau*, which leverages the power of deep-learning splicing prediction to optimise vectors, ensuring they are spliced in the desired manner. Compared to using high-throughput screens to find sequences with desired splicing characteristics (*27*), use of *SpliceNouveau* means only a handful need to be tested. The benefits of the *SpliceNouveau* approach are also apparent from the first iteration in Figure 2C, which demonstrates that naive procedural generation produces sequences that are not predicted to be spliced in the desired manner. Using *SpliceNouveau*, we successfully generated cryptic cassette exons, and also cryptic alternative splice sites, by computationally designing competitor alternative splice sites that are used preferentially when TDP-43 is present. The flexibility and high success-rate of *SpliceNouveau*-designed vectors therefore represents a major step forward in splicing-regulated vector technology (*25*, *27*).

Overall, this study demonstrates how cryptic splicing, ostensibly a driver of ALS and FTD progression, can be reverse-engineered and exploited to enable exceptionally targeted therapeutic protein expression. This approach, which is compatible with conventional AAVs that have previously been approved for gene therapy, can readily be adapted for different transgenes, including tuning of their maximal expression levels and sensitivities to TDP-43 depletion. This will minimise the risk of toxic side-effects and thus improve the chance of obtaining measurable improvements during clinical trials, reducing danger for patients, and increasing incentives for drug developers.

## Acknowledgments

We thank Molly Strom and Ana Cunha of the Crick Institute viral vector core for producing the AAVs, Rupert Faraway for useful discussions on prime editing, Flora C.Y. Lee, Charlotte Capitanchik and Weaverly Lee for feedback on the manuscript, and Sami Barmada for suggestions on validation methods.

## Author contributions

Conceptualization: PF, OGW Software: OGW

Formal analysis: OGW, MZYJC, JJW, ALB Resources: RK, EF, TC, JU, CLP, PF

Supervision: OGW, RK, EMCF, TJC, JU, CLP, PF

Funding acquisition: EMCF, RK, EF, TJC, JU, CLP, PF

Investigation: OGW, MZYJC, JJW, MP, DT, HD, RLS, JD, MEK, MZ, ALB, PRM, PH

Visualisation: OGW, JJW, ALB Methodology: OGW, JJW, CLP, PF Writing - original draft: OGW, PF

Writing - review and editing: MZYJC, JU, CLP, JJW, EMCF

Competing interests: PF and OGW have filed a patent application relating to TDP-REG technology. PF, ALB, MJK and OGW have filed a patent application relating to the use of ASO therapies for correcting cryptic splicing in UNC13A.

## Data and materials availability

RNA-Seq Data for i3Neurons, SH-SY5Y and SK-N-BE(2) are available through the European Nucleotide Archive (ENA) under accession PRJEB42763. Public data was obtained from Gene Expression Omnibus (GEO): iPSC MNs (Klim et al., 2019)-GSE121569, Appocher SK-N-BE(2)-GSE97262, and HeLA Ferguson-GSE136366. NYGC ALS Consortium RNA-seq: RNA-Seq data generated through the NYGC ALS Consortium in this study can be accessed via the NCBI’s GEO database (GEO GSE137810, GSE124439, GSE116622, and GSE153960). All RNA-Seq data generated by the NYGC ALS Consortium are made immediately available to all members of the Consortium and with other consortia with whom we have a reciprocal sharing arrangement. To request immediate access to new and ongoing data generated by the NYGC ALS Consortium and for samples provided through the Target ALS Postmortem Core, complete a genetic data request form at.

## Supplementary Materials

### Materials and Methods

#### Cell culture

SK-N-BE(2) and SH-SY5Y cells were grown in DMEM/F12 (Thermo Fisher Scientific) with 10% FBS (Gibco; Thermo Fisher Scientific). HEK293T cells were grown in DMEM Glutamax (Thermo Fisher Scientific) with 10% FBS (Gibco; Thermo Fisher Scientific).

Transfections were performed with Lipofectamine 3000 (Thermo Fisher Scientific), using 20 μl of Lipofectamine and 20 μl of P3000 reagent per microgram of DNA diluted in Opti-Mem (Thermo Fisher Scientific), following the manufacturer protocol.

A clonal SK-N-BE(2) line expressing a doxycycline-inducible shRNA against TDP-43 was generated by transducing cells with SmartVector lentivirus (V3IHSHEG_6494503), followed by selection with puromycin (1 μg ml−1) for one week.

Polyclonal piggyBac lines were generated by co-transfecting the relevant piggybac vector (backbone from Addgene plasmid #175271) with a vector expressing hyperactive piggyBac transposase (*28*). A 3:1 ratio of transposase vector to expression vector was used. Selection was performed for at least two weeks in 10 μg/ml blasticidin; a control transfection without the transposase expression vector was performed in parallel, to ensure total cell death of transiently transfected cells after selection.

#### Western blotting

Adherent cells were washed with PBS, then lysed in RIPA buffer (25 mM Tris-HCI Buffer pH 7.5, 150 mM NaCl, 1% NP-40, 1% sodium deoxycholate, 0.1% sodium dodecyl sulphate). DNA was sheared via sonication using a Bioruptor Pico device. Samples were loaded onto NuPAGE 4-12% Bis-tris gels (Thermo Fisher Scientific) and transferred to a methanol-activated PVDF membrane using a Mini Trans-Blot Cell (BioRad). Membranes were blocked in 5% fat-free powdered milk in PBS-T buffer (0.2% Tween-20). Primary and secondary incubations were 90 min at room temperature, or overnight at 4°C. Chemiluminescence signal was detected by adding HRP (horseradish peroxidase) substrate. All antibodies used are listed in Supplementary Table S1.

For His-tag pulldown, the Dynabeads His-Tag Isolation and Pulldown (Thermo Fisher Scientific, 10103D) was used as per the manufacturer’s instructions. Cells were lysed using lysis buffer (50 mM sodium phosphate pH 8.0, 1% Triton X-100, 50 mM NaCl in distilled water with cOmplete EDTA-free protease inhibitor (Roche)), then combined with an equal volume 2X Binding/Wash Buffer (50 mM sodium phosphate pH 8.0, 600 mM NaCl and 0.02% Tween-20 in distilled water). Following this, the solution was mixed with Dynabeads magnetic beads and incubated at room temperature for 5 min. After placing the sample on a magnet for 2 min, the supernatant was aspirated and discarded. Subsequently, four washes were performed using 1X Binding/Wash Buffer (diluted to 1x with distilled water), ensuring each wash included thorough resuspension of the beads. Finally, the protein was eluted with His-Elution Buffer (300 mM imidazole, 50 mM sodium phosphate pH 8.0, 300 mM NaCl and 0.01% Tween-20 in distilled water). The eluted sample was mixed with NuPage LDS Sample buffer (4x) (Thermo Fisher Scientific) and the western blot was performed as described above.

#### Cloning

dsDNA fragments were ordered from IDT as GBlocks or EBlocks. PCRs were performed using high-fidelity DNA polymerases (Phusion HF 2x Master Mix or Q5 2x Master Mix; NEB). Plasmid backbones were linearised either via inverse PCR or restriction enzyme digestion. Gibson assembly was performed with 2x HiFi Assembly Master Mix (NEB). Transformation of DNA was performed with Stbl3 bacteria (ThermoFisher Scientific); for the transformation of Gibson assembly products, the reaction mixtures were first purified using SPRI beads to avoid toxicity (Mag-bind TotalPure NGS; Omega-Bio-tek). Ligations were performed using T4 DNA ligase (NEB), following phosphorylation with T4 PNK kinase (NEB). All PCR products that used a plasmid as a template were treated with DpnI (NEB) before downstream steps. All relevant sequences were confirmed either by Sanger sequencing (Source Bioscience or Genewiz) or Nanopore sequencing (Plasmidsaurus).

pPB-EF1a-MegaGate-DD-Blast, which was used as the backbone for piggyBac vectors, was a gift from George Church (Addgene plasmid # 175271; http://n2t.net/addgene:175271; RRID:Addgene_175271) (*29*). pCMV-PEmax, which was used as the basis for CE-containing Prime Editing vectors, was a gift from David Liu (Addgene plasmid # 174820; http://n2t.net/addgene:174820; RRID:Addgene_174820) (*22*); a Tri-Flag-Tagged version was generated via Gibson assembly. pU6-tevopreq1-GG-acceptor, which was used for cloning pegRNAs, was a gift from David Liu (Addgene plasmid # 174038; http://n2t.net/addgene:174038; RRID:Addgene_174038) (*30*). A modified version of the 12QN plasmid was generated via Gibson assembly with the same amino acid sequence as published (*18*).

#### Cell imaging and quantification

Cells were prepared as described above. 96 well plates were seeded with 10,000 cells per well, then transfected as described above using 100 ng of DNA per well. After 52 hours, cells were imaged using an Incucyte microscope (Sartorius), with 20x magnification and 4 images per well. Images were then analysed using CellProfiler (*31*): briefly, red objects were identified using an adaptive threshold (“Robust Background” method), then the total intensity of red signal within these objects was calculated. The mean and standard deviation of the four images for each well, followed by the means of each condition (i.e. each construct +/- doxycycline) were calculated, and the ratio +/- doxycycline was calculated.

#### Quantification of cryptic *AARS1* expression in published RNA-sequencing

Publicly available cell line data were aligned using the pipeline described in (*7*) - briefly, samples were aligned to the GRCh38 genome build using STAR (v2.7.0f) (*32*) with gene models from GENCODE v31 (*33*). NYGC ALS consortium RNA-seq data were processed and categorised according to TDP-43 proteinopathy as previously described (*7*, *34*, *35*). For quantifying the PSI of the *AARS1* cryptic exon, we extracted the counts from the STAR splice junction output tables using bedtools (*36*) for spliced reads mapping to the following coordinates:

chr16 70272882 70276486 AARS1_novel_acceptor-

chr16 70271972 70272796 AARS1_novel_donor-

chr16 70271972 70276486 AARS1_annotated-

We calculated the percent spliced in (PSI) as:

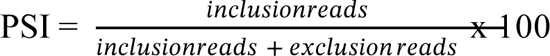

#### SpliceNouveau algorithm

Briefly, the algorithm takes a number of parameters as input; as a minimum, the amino acid sequence to be encoded, and the type and position of the cryptic exon/cryptic splice site, are required. The algorithm then initialises a coding sequence for the given amino acid sequence (or uses an initial sequence provided by the user), and uses SpliceAI (*15*) to assess its predicted splicing behaviour, which is compared to the desired, “ideal” set of splicing predictions. Mutated versions (mutations within the coding sequence are always synonymous) of the sequence are then created and their splicing predictions are calculated. The mutant sequences with the highest “fitness” (i.e. the sequence with a splicing predictions most closely matching the ideal sequence) are retained and used as the basis for subsequent rounds of *in silico* mutagenesis. As such, the process resembles a “directed evolution” approach, but is performed *in silico*. For single intron designs (excluding IR), a competitor splice site is also generated, at a suitable position determined by the algorithm. Additionally, for single introns with alternative 3’ splice sites, the coding sequence upstream of the competitor can be automatically modified with synonymous codons to create a high density of pyrimidines, forming an alternative polypyrimidine tract within the coding sequence, to help generate the competitor splice site.

#### Screening of synonymous Cas9-encoding variants

A long ssDNA oligo containing degenerate bases at the relevant codon wobble positions was ordered from IDT as an Ultramer (“cas9_ultramer”). This oligo was then converted to dsDNA via PCR. The *AARS1*-based reporter was linearised via inverse PCR, deleting the region corresponding to the cas9_ultramer sequence. The two PCR products were combined with Gibson assembly. The Gibson assembly product was purified using SPRI beads and the whole region relevant to splicing (i.e., the candidate CE, its flanking introns, and their flanking exonic sequences) was then amplified via PCR. In parallel, pTwist-CMV was linearised using a primer containing a random barcode. The barcoded linearised vector and PCR-amplified library of candidate CE sequences was then combined via Gibson assembly (HiFi Assembly Master Mix; NEB). Following purification with SPRI beads, the mixture was transformed into Stbl3 bacteria (Thermo-Fisher Scientific), which, following recovery, was transferred directly to ampicillin-LB (without a plating step).

Cas9_ultramer: 5’-GTGTGTGTGTCACCCAGRCTNTCNCGNAARCTNATHAAYGGNATHCGNGAYAARCARTCNGGNA ARACNATHCTNGAYTTYCTNAARTCNGAYGGNTTYGCNAAYCGNAAYTTYATGCARCTNATHCAYGAYGAYTCNCTNACNTTYAARGARGAYATHCARAARGCNCARGTATGCATCACCCCC-3’

The library of plasmids was purified, then transfected into SK-N-BE(2) cells with or without doxycycline-inducible TDP-43 knockdown. RNA was purified reverse transcribed using a reverse transcription primer featuring a UMI (unique molecular identifier); this was then amplified via PCR using primers with Illumina-compatible overhangs. The resulting PCR products were sequenced via Illumina sequencing. The reads were analysed using custom python and R scripts (available at https://github.com/Delayed-Gitification/TDP-REG-Paper).

#### Nanopore analysis

Targeted RT-PCR was performed using vector-specific primers. Barcoding was performed either using custom barcoded primers, or via barcode ligation with kit SQK-NBD114.24. R9.4.1 flow cells were used for all experiments except those for measuring prime editing levels, for which R10.4.1 flow cells were used. Basecalling was performed using Guppy v6.0.1, with the relevant “SUP” (super-accuracy) model. Demultiplexing was performed using the demultiplex_nanopore_barcodes.py function from nano_tools v0.0.1 (https://github.com/Delayed-Gitification/nano_tools). Alignment was performed using Minimap2 (v2.1) (*37*). Pileups were generated using the perform_enhanced_pileup.py function from nano_tools v0.0.1. Splicing analysis was performed by extracting splice junctions from reads using the extract_splice_junctions_from_bam.py function from nano_tools v0.0.1, followed by analysis with custom R scripts (available in https://github.com/Delayed-Gitification/TDP-REG-Paper).

#### Generation of AAVs

Vectors were generated using Gibson assembly and full nanopore sequencing (Plasmidsaurus) was used to validate the sequences; all vectors featured the human Synapsin promoter. rAAVs were produced by triple transduction of HEK-293T cells essentially as described in Challis et al. (*38*), with the following exceptions: 2-4 15 cm dishes were used and after the PEG precipitation samples were resuspended in 4 ml, 3 ml chloroform was added, mixed for 2 min by vortexing, and centrifuged at 3000 g for 20 min and the aqueous layer was loaded on iodixanol gradients in a Type 70.1 rotor and centrifuged for 2 h at 52000 RPM. The rAAV sample was collected and buffer exchanged with 1x PBS 5% Sorbitol 0.1 M NaCl (0.25 M NaCl final). Addgene plasmid #103005 was used for AAV production (*39*). Titres were between 1.0E14 and 3.1E14 genome copies per ml.

#### Mice

All animal care and experimental procedures were performed in accordance with animal study proposal ASP23-003 approved by the National Institute of Child Health and Human Development Animal Care and Use Committee. TDP-43^Fl/wt^ (*Tardbp^tm1.1Pcw^*/J) mice obtained from Dr. Philip Wong at Johns Hopkins University (Jax stock No. 017591) were crossed to homozygosity then crossed to the Chat-IRES-Cre::deltaNeo line (***Chat^tm1(cre)Lowl^***/J);Jax Stock No. 031661), in which the neomycin resistance cassette was removed to avoid ectopic expression sometimes observed in the ChAT-IRES-Cre line. This produced male TDP-43^Fl/wt^;Chat-Cre^+/+^ breeders which were crossed to female TDP-43^Fl/Fl^ mice to generate both TDP-43^Fl/Fl^;Chat-Cre^+/wt^ and TDP-43^Fl/wt^;Chat-Cre^+/wt^ that were used in these experiments. The positive control mScarlet AAV was injected into mice with a Sun1-tag (TDP43 fl/fl;Sun1-GFP +/+), which were generated by crossing the previously described TDP43 fl/fl mice to CAG-Sun1/sfGFP mice (B6;129-*Gt(ROSA)26Sor^tm5(CAG-Sun1/sfGFP)Nat^*/J; StockNo: 021039). The genetic background was C57Bl/6J for all animals. All animal ages are listed in Supplementary Table S2.

#### Intracerebroventricular Injections

A 10 μl Hamilton syringe (65460–06) with a 33G needle (65461-02) was loaded with up to 10 μl of undiluted virus and placed on a syringe pump (KD Scientific 78-0220). p0-p2 pups were anaesthetised on ice for approximately 1 minute. After anaesthesia, pups were placed on a sterilised mobile surface and moved into the Hamilton syringe. 1 μl of virus was delivered at 1 μl/min into the left ventricle, approximately 1 mm lateral from the sagittal suture. The syringe was kept in place for approximately 30 seconds after the injection before being removed to reduce backflow. Pups were placed on a heating pad to recover before being returned to their dam in the home cage. Injection details are in Supplementary Table S2.

#### Tissue Preparation and Sectioning

Mice were anaesthetised (age 3-7 weeks) with I.P. injections of 2.5% avertin and transcardially perfused with 10mL of 1X PBS followed by 10mL of 4% paraformaldehyde. Spinal cords were dissected and placed in 4% paraformaldehyde overnight before being placed in a 30% sucrose in PBS cryopreservation solution. After 24 hrs in cryopreservation solution, lumbar spinal cords were embedded in O.C.T (Tissue-Tek) and frozen. Frozen blocks were sectioned into 16 μm-thick coronal slices onto positively charged slides using a Leica CM3050 S Research S Cryostat. Slides were stored at −80°C for up to 2 weeks.

#### Immunostaining of tissue

Slides were removed from −80°C and thawed to room temperature, then washed in 1X PBS before being placed in citrate buffer (pH 6.0) for antigen retrieval. The solution and slides were microwaved for 45 seconds (until light boil) and allowed to cool back to room temperature. Tissue was then permeabilized in 0.1% Triton-X100 in 1X PBS (PBSTx), then blocked in 5% normal donkey serum in 0.1% PBSTx. Primary antibodies were diluted in 0.5% normal donkey serum in 0.1% PBSTx and incubated overnight at 4°C. Slides were then washed in 0.1% PBSTx and incubated for 1 hr in secondary antibody (ThermoFisher) diluted in 0.1% PBSTx. After a final 1X PBS wash, slides were coverslipped with Prolong Diamond (ThermoFisher P36961) and dried overnight at room temperature in the dark before being stored at 4°C. Primary antibodies: Rat anti-TDP-43 (Biolegend #808301, 1:3000), Rabbit anti-RFP (Rockland #600-401-379, 1:100), Guinea Pig anti-VAChT (Synaptic Systems #139105, 1:500).

#### Imaging and Analysis

Up to 6 slides were loaded simultaneously onto the Olympus VS200 slide scanner for imaging. Slides were imaged at 20X with 5 z planes, 2 µm apart with the following filer cubes: DAPI, FITC, TRITC, and Cy5. For motor neuron counts, maximum intensity projections were used. Briefly, ROIs were drawn around motor neurons based on VAChT and DAPI by an investigator blind to genotype. To determine RFP positive vs negative motor neurons, the brightness was adjusted so that no RFP was detectable in background spinal cord regions or in between motor neurons. Any motor neurons that still had detectable RFP were considered positive while those with background levels were considered negative.

#### Cytoplasmic aggregation

A plasmid encoding SNAP-TDP-43-12QN with an L207P mutation in the human TDP-43 sequence was generated; the sequence of the TDP-43 and 12xQN repeat was identical to a published study (*18*). TDP-REG mScarlet reporter constructs were co-transfected with either SNAP-TDP-43-12QN or pHTN HaloTag® CMV-neo Vector (with no insert; Promega) at a 1:3 mass ratio of reporter:TDP/Halo plasmid.

For confocal imaging, cells were plated on Matrigel-coated Ibidi 8-well microscopy dishes, then transfected after 24 hours. Three hours prior to fixation, SNAP-Cell Oregon Green (NEB) prepared in DMSO (in accordance with the manufacturer recommendations) was diluted into growth media (1:2,000 v/v) and the original media was replaced with this media. 60 min prior to fixation, the SNAP-Cell Oregon Green media was replaced with normal growth media. HEK293T cells were fixed 28 hours after transfection, whereas SK-N-BE(2) cells were fixed 50 hours after transfection, to compensate for their lower protein expression levels; fixation was performed in 2% PFA (methanol-free) diluted in PBS for 20 min at room temperature. Cells were permeabilized with 0.1% Triton X100 diluted in PBS for 10 min at room temperature. Blocking was performed for one hour with 0.1% Tween 20 and 2% W/V bovine serum albumin (BSA) diluted in PBS. Primary antibody (TDP-43 antibody; Proteintech) was diluted 1:500 into PBS with 0.1% Tween 20 and cells were incubated overnight at 4°C. After three washes with PBS-Tween 20, secondary antibody (anti-Rabbit IgG conjugated to Alexa-fluor 647; Abcam) was diluted into PBS with 5% w/v BSA and 0.05% Tween 20, and incubation was performed for 60 min at room temperature. Finally, samples were washed and incubated with DAPI for nuclear staining. Imaging was performed on a Zeiss 880 Inverted Confocal using a 63x oil objective lens. To reduce bleed-through from the far-red channel into the red channel, a narrow wavelength cut-off was used for the native mScarlet signal of approximately 575-595 nm.

For imaging with the Incucyte, HEK293T cells were plated into a 96 well plate. The following day, they were transfected with 70 ng of SNAP-12QN-TDP-43 plasmid or empty Halo-tag vector, 20 ng of mScarlet reporter plasmid and 10 ng of mGreenLantern plasmid (as a transfection control). Media was changed 24 hours after transfection. Imaging on an Incucyte S5 was performed 48 hours after transfection.

#### Prime editing

*SpliceNouveau* was used to design a cryptic exon within the pCMV-PEMax vector. The vector was transfected in SK-N-BE(2) cells with or without TDP-43 knockdown and the splicing of the construct was analysed via RT-PCR. Primers are listed in Table S3.

The PrimeDesign web tool was used to design the pegRNA and nicking sgRNA (*40*).

The pegRNA used had sequence 5’-GTAAAAGCATGGATGGAGAGAGTTTTAGAGCTAGAAATAGCAAGTTAAAATAAGGCTAGTCCGTTATCAACTTGAAAAAGTGGCACCGAGTCGGTGCATGGACTCACGCATCTCTCCATCCATGCCGC GGTTCTATCTAGTTACGCGTTAAACCAACTAGAATTTTTTT-3’

The sgRNA used had sequence 5’-GAAACACCGTGGGGATAAGAGTTCTTTCCGTTTTAGAGCTAGAAATAGCAAGTTAAAATAAGGCTAGTCCGTTATCAACTTGAAAAAGTGGCACCGAGTCGGTGCTTTTTT-3’ pCMV-PEMax or a version containing a cryptic exon were transfected into SK-N-BE(2) cells with or without doxycycline-inducible TDP-43 knockdown. 600 ng of prime editing vector was used, in addition to 200 ng of pegRNA, 100 ng of nicking sgRNA and 100 ng of a plasmid expressing mScarlet and the blasticidin resistance gene. 24 hours after transfection, the media was changed and supplemented with 10 µg/ml blasticidin to select for transfected cells. After an additional 48 hours, the cells were subcultured, and after six days total the samples were harvested and gDNA was purified. gDNA was amplified via PCR and amplicons were analysed by Nanopore sequencing. Pileups were generated using the perform_enhanced_pileup.py function from nano_tools v0.0.1 (https://github.com/Delayed-Gitification/nano_tools). The fraction of reads containing the expected edit was calculated.

#### Luciferase analysis

A modified Gaussia luciferase amino acid sequence, in which the methionines are replaced to improve resistance to oxidation, was reverse translated and optimised by SpliceNouveau, including internal cryptic exons. Vectors were transfected into SK-N-BE(2) cells with or without doxycycline-inducible TDP-43 knockdown. Luciferase activity was measured by extracting an aliquot of cell media and mixing with Pierce™ Gaussia Luciferase Glow Assay Kit (Thermo Fisher Scientific) following the manufacturer protocol. Splicing was analysed via targeted Nanopore sequencing, using the approach described above.

#### Design of TDP-43/Raver1 expression vectors

The TDP-43/Raver1 protein sequence used for all experiments was: MG<u>PKKKRKV</u>EDPGG*<u>PAAKRVKLD</u>*GG**YPYDVPDYA**GGM**<u>SEYIRVTEDENDEPIEIPSEDDGTVLLSTVTAQFPGACGL</u> <u>RYRNPVSQCMRGVRLVEGILHAPDAGWGNLVYVVNYPKDNKRKMDETDASSAVKVKRAVQKTSDLIVLGLPWKTTEQ</u> <u>DLKEYFSTFGEVLMVQVKKDLKTGHSKGFGFVRFTEYETQVKVMSQRHMIDGRWCDCKLPNSKQSQDEPLRSRKVFV</u> <u>GRCTEDMTEDELREFFSQYGDVMDVFIPKPFRAFAFVTFADDQIAQSLCGEDLIIKGISVHISNAEPKHNSN</u>***LPPLL GPSGGDREPMGLGPPATQLTPPPAPVGLRGSNHRGLPKDSGPLPTPPGVSLLGEPPKDYRIPLNPYLNLHSLLPSSNLAGKETRGWGGSGRGRRPAEPPLPSPAVPGGGSGSNNGNKAFQMKSRLLSPIASNRLPPEPGLPDSYGFDYPTDVGPRRLFSHPREPTLGAHGPSRHKMSPPPSSFNEPRSGGGSGGPLSHF*\*

In the above sequence, the SV40 NLS is underlined, the c-Myc NLS is underlined and in italics, the HA-tag sequence is in bold, the TDP-43 sequence is in bold and underlined, and the C-terminal Raver1 sequence is in italics. The 2FL mutation changed the sequence HSKGFGF within RRM1 to HSKGLGL.

*SpliceNouveau* was used to design constructs with internal cryptic exons within the region encoding TDP-43 within the TDP-43/Raver1 sequence above. For initial screening of their cryptic exon properties, the designed sequences were cloned into a mammalian expression vector (pTwist-CMV) containing the 2FL mutation. These were then transiently transfected into SK-N-BE(2) cells with or without doxycycline-induced TDP-43 knockdown. RNA was extracted, reverse transcribed and RT-PCRs were performed to analyse inclusion of the synthetic TDP-43-encoding cryptic exons (primers listed in Table S3), which was aided by the use of a QIAxcel Advanced machine (QIAGEN).

#### Rescue of endogenous cryptic splicing with TDP-43/Raver1

The piggyBac system was used to make SK-N-BE(2) cell lines with constitutive EF1A promoters driving expression of constitutive (i.e. without a cryptic exon) or two cryptic exon-containing TDP-43/Raver1 constructs, or mScarlet; a PGK promoter drove expression of the blasticidin resistance gene (*29*). Note that these lines were made using the clonal line that featured the doxycycline-inducible TDP-43 shRNA, enabling doxycycline-inducible TDP-43 knockdown. Polyclonal lines were produced in triplicate. Each polyclonal line was then plated with or without doxycycline-treatment (1000 ng/ml) for six days, then harvested for western blotting or RT-PCRs. Western blotting was performed as described above.

RT-PCRs for *UNC13A*, *STMN2* and *AARS1* were performed via reverse transcription with Superscript IV (Thermo Fisher Scientific) followed by PCR using either Q5 2x Master mix (NEB) or One-Taq Quickload 2x Master Mix (NEB). *UNC13A* and *AARS1* PCRs used two primers, whereas *STMN2* used two reverse primers because the STMN2 cryptic exon induces premature polyadenylation; primer sequences are listed in Table S3. PCR products were electrophoresed on a QIAxcel advanced system. Raw data was exported and analysed in R using the QIAxcelR package (v0.1) (https://github.com/Delayed-Gitification/QIAxcelR; see figures markdown for analysis code used).

#### Growth competition assay

The piggyBac system was used to generate SK-N-BE(2) cells with stably-integrated, doxycycline-inducible expression of TDP-43/Raver1 fusion, either constitutive or with an internal cryptic exon, or mTag-BFP2 blue fluorescent protein. Cell lines were generated in triplicate. 100 ng of each vector, plus 400 ng of hyperactive piggyBac transposase, were used per 24 well transfection. Note that in this case, the SK-N-BE(2) cells did not feature the doxycycline-inducible TDP-43 shRNA cassette.

Following selection of stable cells in 10 µg/ml blasticidin for 20 days, blasticidin was removed and the different cell lines were mixed together, again in triplicate. Each mix was placed into three different wells, with 0, 30 or 1000 ng/μl of doxycycline. After three days, the cells were subpassaged. After a further seven days, the cells were lysed in 20 mM Tris-HCl pH 7.5, 0.5 mM EDTA, 1% Trixon X100 and 500 ng/μl proteinase K; samples were incubated at 55°C for 20 min, then 96°C for 5 min.

A multiplex PCR was performed using two pairs of primers: one pair which amplified the three TDP-43/Raver1 vectors, and another pair which amplified the BFP construct. PCR was performed with Q5 2x Master Mix (NEB), using 2.5 μl of lysate into a 25 μl PCR. Primers are listed in Table S3. The samples were then purified and barcoded Nanopore libraries were prepared using the Native Barcoding 24 kit with R10.4.1 chemistry. The library was sequenced with an R10.4.1 Flongle device and High Accuracy basecalling was used in real-time with Guppy. Reads were then aligned to a “reference genome” consisting of the four constructs with Minimap2 (v2.1) (*37*), and mapping statistics were calculated by analysing the resulting bam files in R.

---

**Fig. S1.**
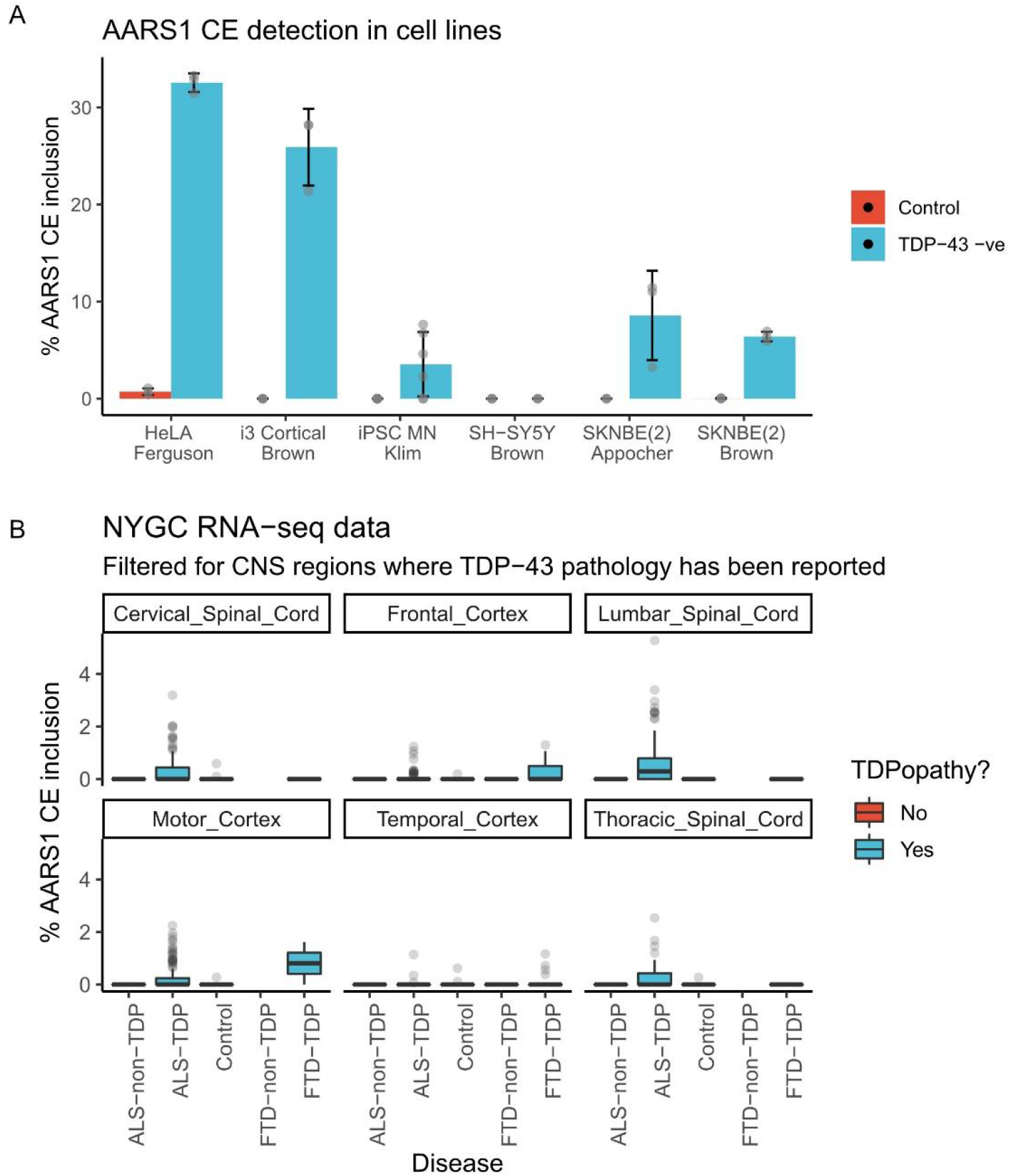
**A:** *AARS1* cryptic exon inclusion percentage for various cell lines in which TDP-43 levels were reduced artificially; error bars show standard deviation. **B:** *AARS1* cryptic exon inclusion for bulk RNA seq from various tissues and patients in the NYGC dataset.

**Fig. S2.**
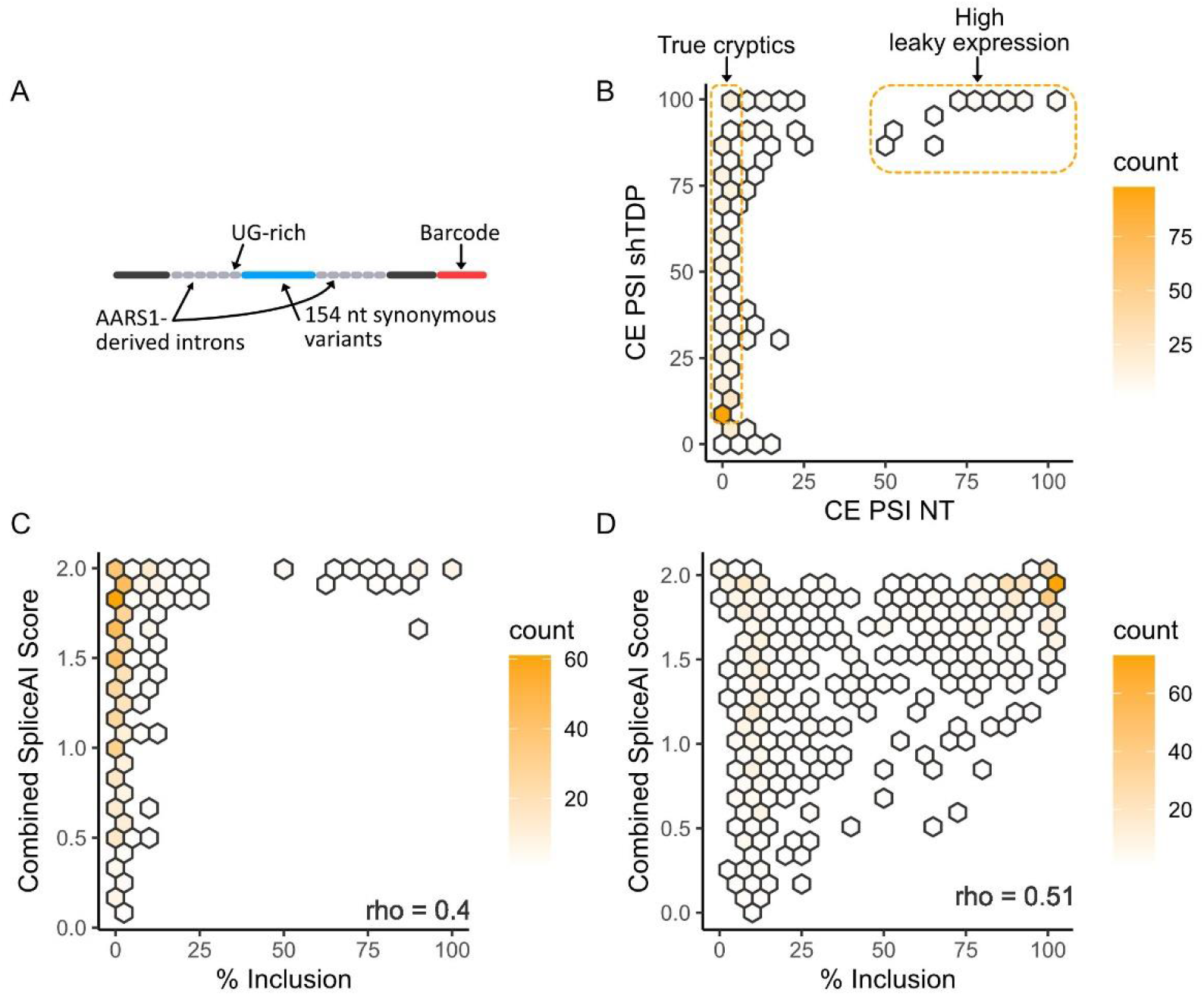
**A**: Schematic of the barcoded library of vectors containing different candidate CE sequences, each encoding the same amino acid sequence (in this case, a fragment of S. pyogenes Cas9) but with different codon optimisation. **B:** Heatmap of % CE inclusion for library of Cas9-fragment-encoding cryptic exons in SK-N-BE(2) cells with TDP-43 knockdown versus untreated cells (“NT” = “not treated”). Areas corresponding to “True cryptics” (i.e. those that are only expressed upon TDP-43 knockdown) and “High leaky expression” (i.e. those that are expressed regardless of TDP-43 knockdown) are highlighted. **C:** Heatmap of the SpliceAI score of the candidate CE acceptor and donor splice sites against the PSI of each CE in SK-N-BE(2) cells without TDP-43 knockdown (i.e. untreated cells); Spearman correlation shown (bottom)**. D:** Heatmap of the SpliceAI score of the candidate CE acceptor and donor splice sites against the PSI of each CE in SK-N-BE(2) cells with TDP-43 knockdown; Spearman correlation shown.

**Fig. S3.**
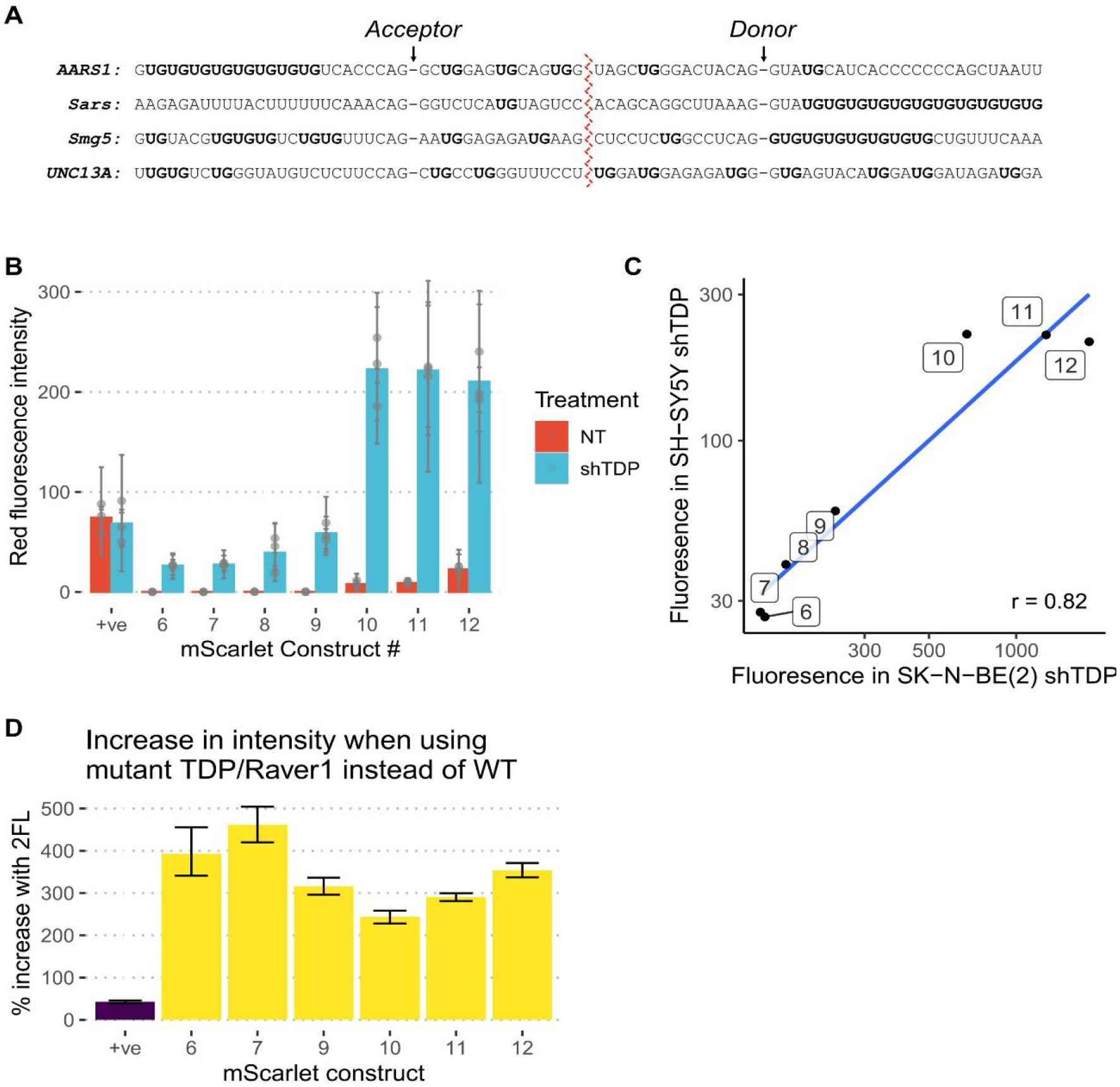
**A:** Diagrams of four endogenous CEs, highlighting the position of UG dinucleotides relative to the splice sites. **B:** Quantification of mScarlet fluorescence from SY-SY5Y cells transfected with mScarlet constructs 6-12; dots show average of each well; error bars show standard deviation within each well. **C:** A comparison of the fluorescence of the seven TDP-REGv2 mScarlet constructs in SY-SY5Y and SK-N-BE(2) cells, with TDP-43 knockdown; Pearson correlation is shown. **D:** A comparison of the fluorescence of six TDP-REGv2 mScarlet constructs and a positive control transfected into SK-N-BE(2) cells with TDP-43 knockdown, when co-transfected with a mutant TDP-43/Raver1 fusion protein (which is unable to rescue splicing) versus when co-transfected with a functional TDP-43/Raver1 fusion protein; error bars show 90% confidence interval as determined by Monte-Carlo methods.

**Fig. S4.**
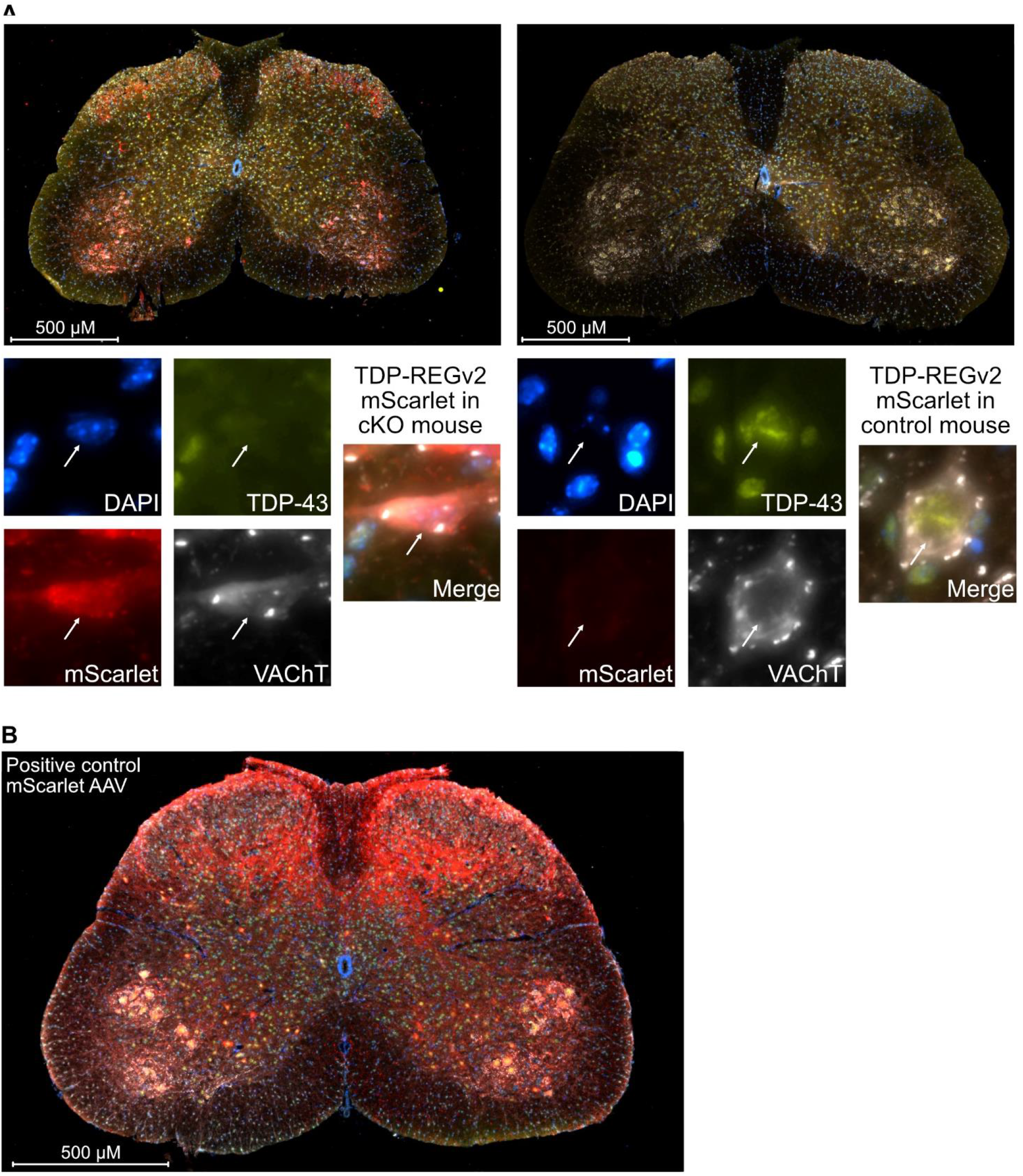
A: Fluorescence microscopy of spinal cords sections from TDP-43 cKO or control mice (left and right respectively) injected with a TDP-REGv2 mScarlet AAV (construct #7). Magnifications of two representative motor neurons are shown below. Blue = DAPI, Yellow = TDP-43, White = VaChT, Red = mScarlet). B: Representative fluorescence microscopy image of a spinal cord section from a control mouse injected with a positive control mScarlet AAV (i.e. without TDP-REG); colouring is the same as Part A.

**Fig. S5.**
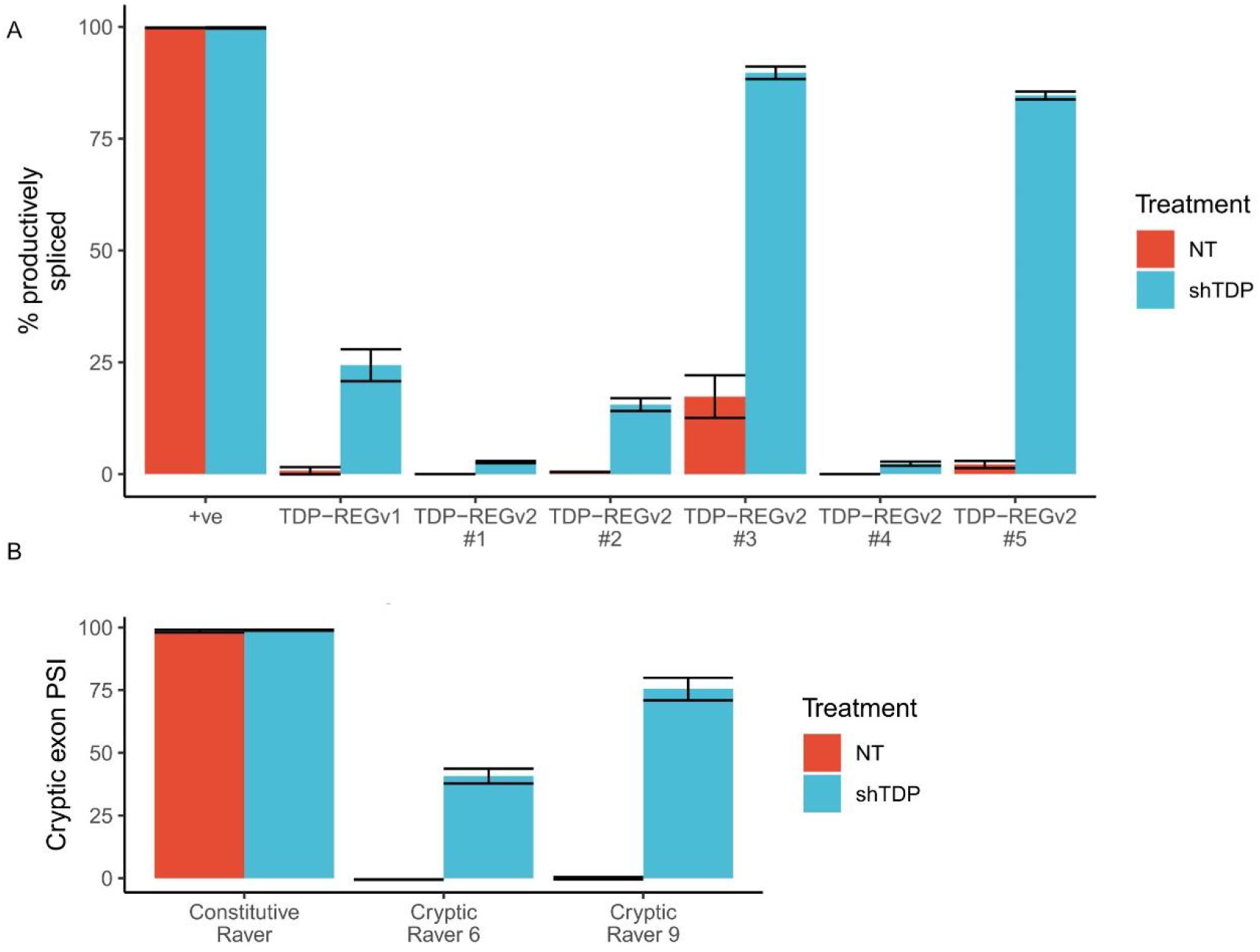
**A:** Nanopore sequencing of seven Luciferase constructs (one positive control, one TDP-REGv1, and five TDP-REGv2) with or without TDP-43 knockdown; error bars show standard deviation across replicates. **B:** Quantification of RT-PCRs detecting the internal cryptic exons present in TDP-REGv2 TDP-43/Raver1 constructs. In contrast with the data shown in Figure 4, in this figure the cells were stably-expressing the vectors, and the TDP-43/Raver1 fusion protein was functional (i.e. without the 2FL mutation); error bars show standard deviation.

**Fig. S6.**
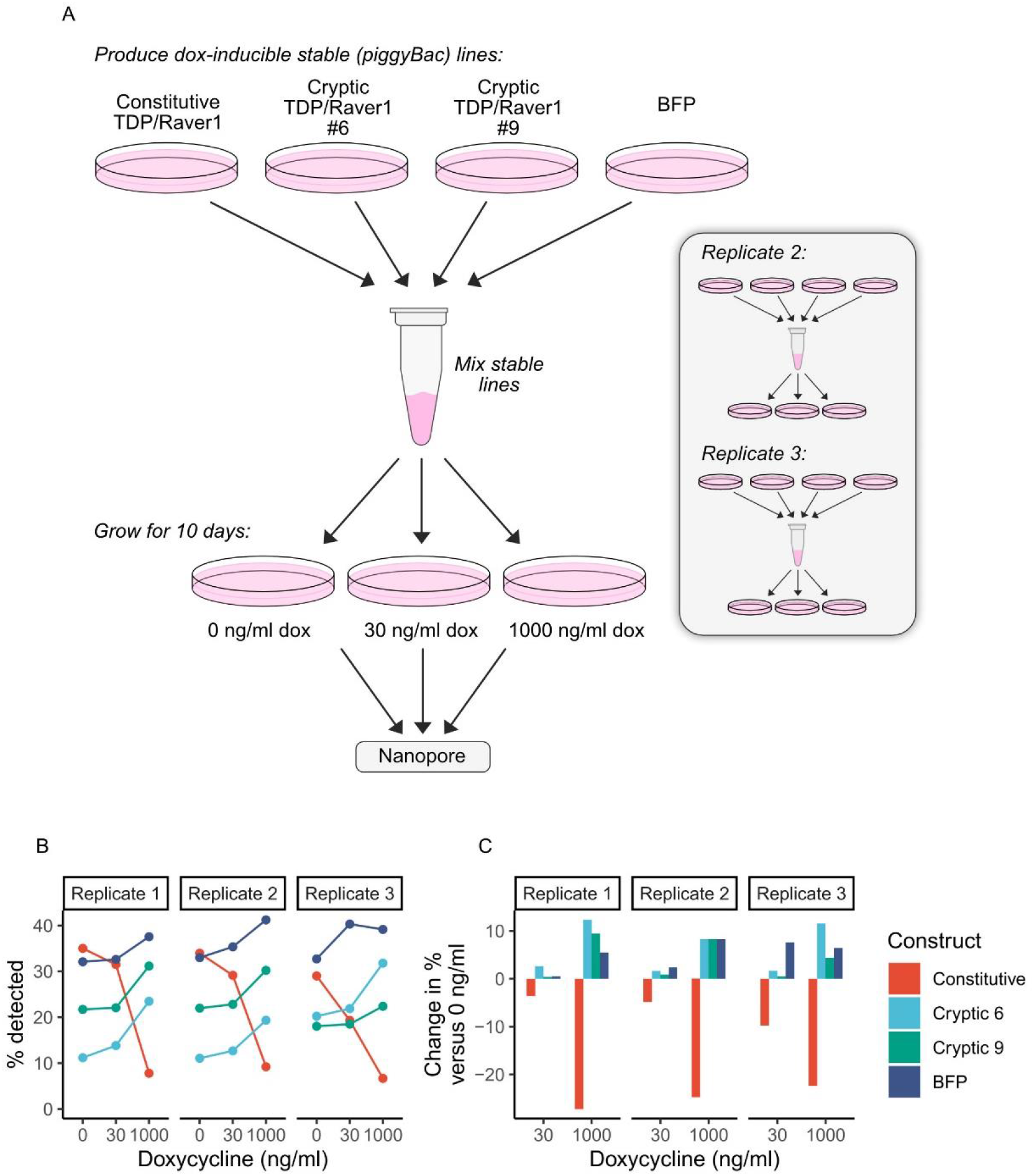
**A:** A schematic showing the experimental procedure for the growth competition assay. **B:** Quantification of Nanopore reads derived from each of the four constructs used to make stable piggyBac lines for each doxycycline concentration and each replicate. (Note that differences in % at 0 ng/ml doxycycline can be explained by PCR bias during Nanopore library preparation and unequal initial mixtures of the different lines; comparisons are only valid between doxycycline concentrations within the same replicate.) **C:** A second visualisation of the same Nanopore data, where values for each construct are compared with their equivalent value when no doxycycline was added, for clarity.

**Table S1.**
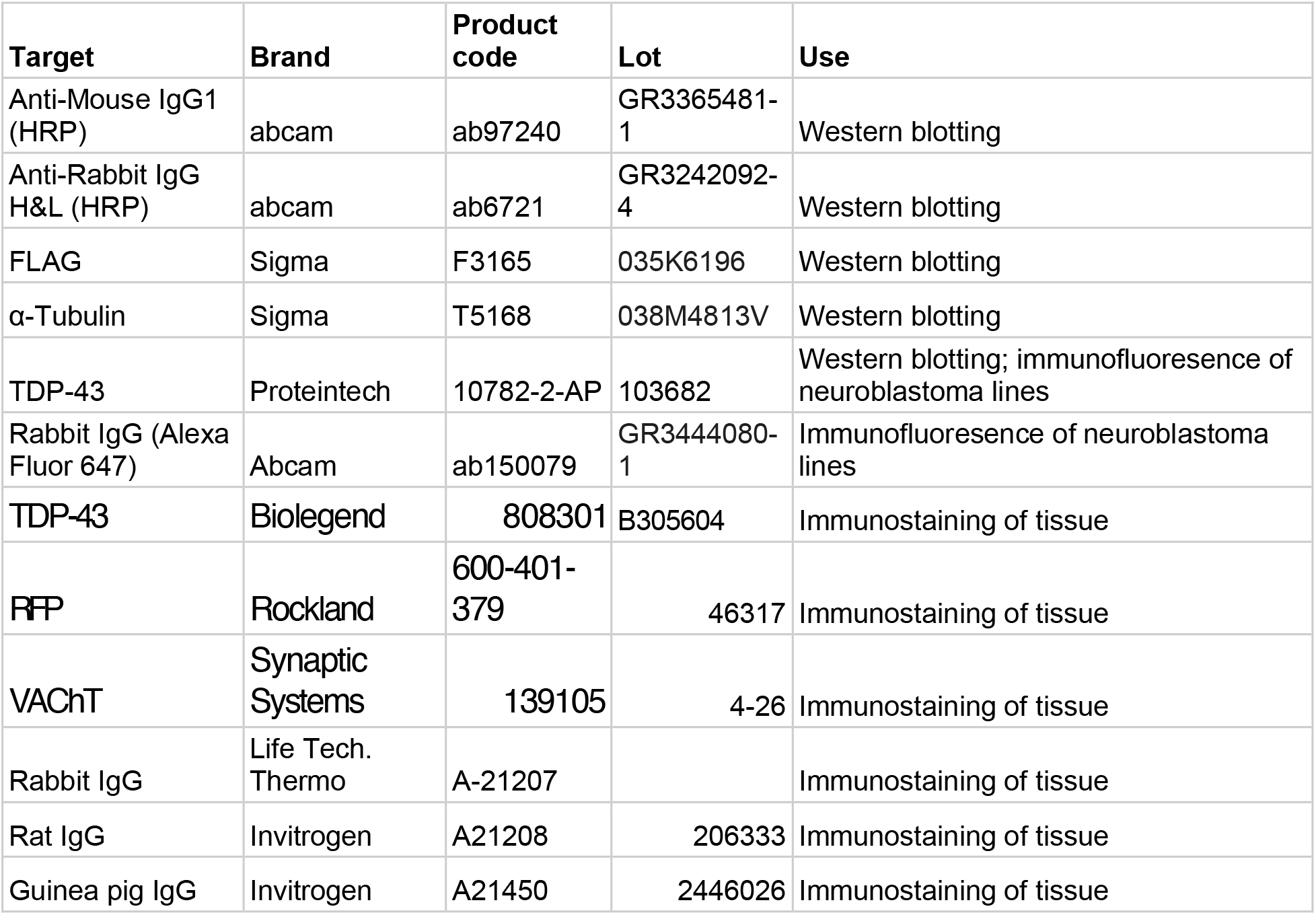
Antibodies used in western blotting and microscopy experiments.

**Table S2.**
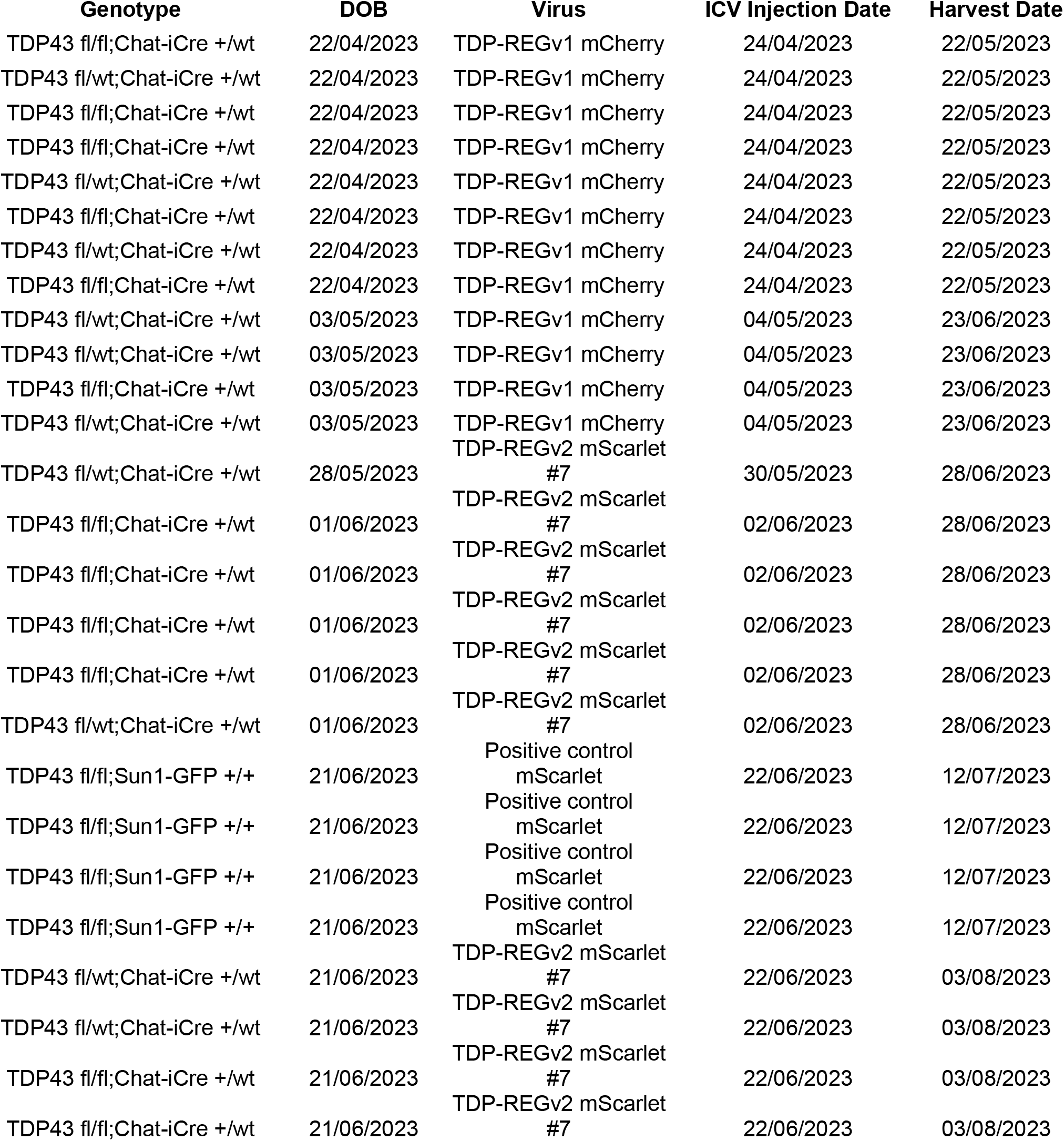
Details of animal experiments.

**Table S3.**
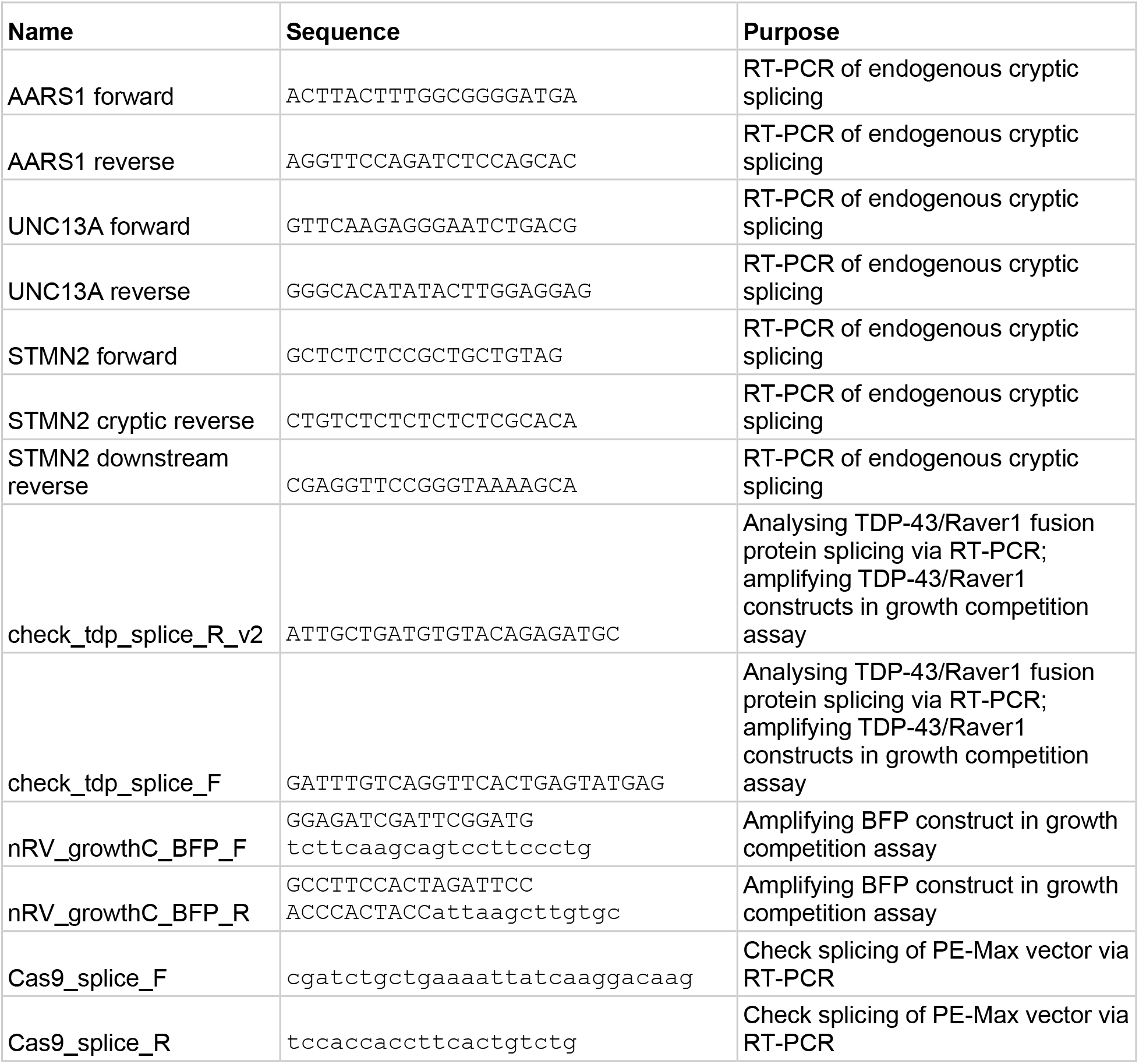
Primers used for RT-PCRs.

